# Connectome lateralization in autism across the first 14 years: heterogeneity related to developmental stage, hemisphere, and pathophysiology

**DOI:** 10.64898/2026.03.02.708940

**Authors:** Qi Liu, Qingyang Li, Xuelian Li, Xinlin Wei, Xiaodong Zhang, Wei Zhou, Lingli Zhang, Tai Ren, Congying Huang, Hangyu Tan, Like Huang, Keyi Liu, Jingyu Chen, Wenjun Xu, Qianlong Zhang, Keith M. Kendrick, Weihua Zhao, Fei Li

## Abstract

Hemispheric lateralization is fundamental feature to neurodevelopment, yet its dynamic evolution in autism across childhood and underlying molecular mechanisms remain unknown. Thus, using a large-scale resting-state fMRI dataset (n = 1553, ages 1-14), we investigated developmental heterogeneity in connectome lateralization. At the group level, abnormalities in dynamic connectome lateralization strength (DCLS) progressed from focal patterns in early childhood (≤ 6 years) to widespread distributions by late childhood (≤ 14 years), independent of symptom severity. The late-stage patterns were validated using the ABIDE dataset (n=153). Late childhood also showed marked increases in inter-individual heterogeneity. These alterations were significantly associated with clinical phenotypes, transcriptomic profiles, and multiple neurotransmitter systems, with distinct patterns across childhood groups. Our findings reveal a developmentally dynamic trajectory of connectome lateralization in autism and highlight late childhood as a critical window for stage- and individual-specific interventions tailored to substantial individual heterogeneity.

## Introduction

Neuroimaging has become an indispensable tool for mapping the trajectory of brain development, quantifying both rapid structural maturation^1–3^ and continuous functional refinement^4,5^ that underpin the emergence of core cognitive, social, and emotional abilities^6,7^. The remarkable neuroplasticity characteristic of childhood (ages 1–14) enables this development but also confers a state of heightened vulnerability. Adverse environmental or biological disruptions during this period can significantly alter typical developmental trajectories, contributing to the onset of neurodevelopmental disorders^8^. Autism spectrum disorder, for instance, typically first manifests during early childhood^9^. Despite its recognized importance, the core neuropathological mechanisms that underlie autism across the entire span of childhood remain poorly understood.

Hemispheric lateralization is a defining feature of human brain organization, with asymmetries in structure, connectivity, and gene expression already detectable as early as in fetuses and neonates, supporting efficient neural processing across cognitive and socio-communicative domains^10–13^. Growing evidence indicates that deviations from typical lateralization are closely associated with neurodevelopmental and psychiatric dysfunction^14^. In individuals with autism, these alterations have been characterized by atypical asymmetries in the morphological and functional features^15,16^, and connectivity of inter-hemispheric homotopic regions^17,18^, as well as the topological properties of intra-hemispheric networks^19,20^. Notably, inter-hemispheric communication is not confined to homotopic connections, as the corpus callosum also mediates extensive inter-hemispheric heterotopic pathways^21^. Coupled with intra-hemispheric circuits, these diverse connections contribute to a balance between inter- and intra-hemispheric connectivity, which is essential for supporting complex cognitive and behavioral functions^22,23^. Aligning with this framework, using resting-state functional magnetic resonance imaging (rsfMRI) data, previous studies have quantified the difference between inter- and intra-hemispheric functional connectivity (FC) for a given brain region to index this intrinsic balance^24,25^. This approach has proven effective in revealing neurobiological mechanisms and potential biomarkers across various neuropsychiatric conditions^24,25^. Nevertheless, traditional approaches on functional hemispheric lateralization have predominantly focused on static connectivity, which assumes temporal stability and is inherently influenced by scanning duration of rsfMRI^26^. Emerging evidence has highlighted the efficacy of dynamic approaches (e.g., dynamic conditional correlations, DCC) for assessing time-varying functional connectivity strength, which has proven highly sensitive in identifying disease-related brain disruptions^27,28^. Remarkably, the alterations in dynamic functional lateralization and their association with autism have remained unexplored. Investigating the dynamics of such lateralization patterns in autism may provide valuable insights into the neural mechanisms underlying their complex clinical phenotypes.

Although a reliable clinical diagnosis of autism can be made as early as 12 months based on DSM-5^29^, understanding the corresponding neural substrates is hindered by a scarcity of neuroimaging data in early childhood (1-6 years). A substantial proportion of earlier studies have relied chiefly on datasets such as the Autism Brain Imaging Data Exchange (ABIDE) dataset that feature children older than six years^30,31^, leaving the developmental trajectories of functional lateralization largely unmapped^17,18^. Moreover, one recent meta study highlights that atypical functional lateralization in autism extends beyond group-level mean differences to manifest as higher inter-individual heterogeneity when compared to typically developing individuals (TD)^16^. Theoretically, regions exhibiting homogeneous effects are indicative of a shared pathophysiological process, while regions demonstrating high variability are suggestive of non-overlapping subtypes^28,32^. However, the association between these mean-level alterations and inter-individual heterogeneity in functional lateralization, and specifically how this relationship is modulated by distinct developmental stages, remains to be characterized in autism. Additionally, transcending these macroscopic characterizations to elucidate the molecular substrates of atypical brain representations is essential for establishing a mechanistic understanding of neurodevelopmental and psychiatric disorders^33,34^. Recent advancements in brain-wide transcriptional resources, such as the Allen Human Brain Atlas^35,36^, together with spatial maps of neurotransmitter transporter and receptor distributions^37^, have established a powerful multimodal framework for linking alterations in functional lateralization to their underlying genetic and neurochemical substrates. However, few studies have delineated how dynamic functional lateralization evolves across the full span of childhood in autism, and the molecular mechanisms that shape these developmental patterns remain unknown.

To bridge these gaps, the present study leveraged a large-scale rsfMRI dataset comprising 1-14 years children (N = 1553, autism=1227), encompassing the critical developmental period from early to late childhood. We combined the DCC approach with a quantitative assessment of inter- and intra-hemispheric functional connectome differences to construct a dynamic connectome lateralization index. By further integrating individual clinical data and transcriptional resources, we systematically disentangled the developmental heterogeneity of dynamic lateralization in autism, aiming to decode how these macroscopic functional anomalies map onto clinical phenotypes and their underlying molecular architecture. Specifically, we aimed to: (1) delineate the spatiotemporal trajectories of dynamic functional lateralization, characterizing how group-level differences and inter-individual heterogeneity evolve from early (1 ≤ age ≤ 6 years) to late childhood (6 < age ≤ 14 years), (2) determine the clinical relevance of this lateralization by examining their associations with individual symptom severity; and (3) elucidate its biological basis by anchoring these macroscopic phenotypes to the spatial distribution of brain-wide gene expression and neurotransmitter receptors.

## Results

### Analytical Framework

To capture the full developmental spectrum from early to late childhood, we included rsfMRI data and clinical measurements from 1,227 autistic and 326 TD children in the Shanghai Autism Early Developmental Cohort (SAED, 1 ≤ age ≤ 14 years, Supplementary Table 1)^15,38^. To enhance replication and validation, we further included rsfMRI data from 56 autistic and 97 TD individuals with ages restricted to 6-14 years from six sites of Autism Brain Imaging Data Exchange Cohort (ABIDE I and II, Supplementary Table 2)^30,31^. Dynamic connectomes were estimated using the DCC approach^28,39,40^, and the dynamic connectome lateralization strength (DCLS) index was subsequently calculated to quantify the degree of lateralization at the hemispheric connectome-scale for each brain region (Fig. 1a). We next characterized developmental-stage–specific (early childhood: 1 ≤ age ≤ 6 years; late childhood: 6 < age ≤ 14 years) and hemispheric patterns of DCLS abnormalities in autistic individuals from the SAED cohort, and tested the reproducibility of the late childhood findings in the ABIDE cohort. Furthermore, by integrating normative modeling with inter-individual heterogeneity analysis^32,41^, we characterized developmental- and hemispheric-specific variations in DCLS heterogeneity among individuals with autism (Fig. 1b). Subsequently, multivariate associations between whole-brain DCLS values and clinical measurements were examined using sparse canonical correlation analysis (sCCA, Fig. 1c)^42–44^. Finally, by integrating transcriptomic profiles from the Allen Human Brain Atlas (AHBA, http://human.brain-map.org)^35,36^ and positron emission tomography (PET)–derived neurotransmitter maps from over 1,200 healthy participants^37^, we further investigated the molecular and neurochemical substrates underlying DCLS abnormalities (Fig. 1d).

**Fig. 1.**
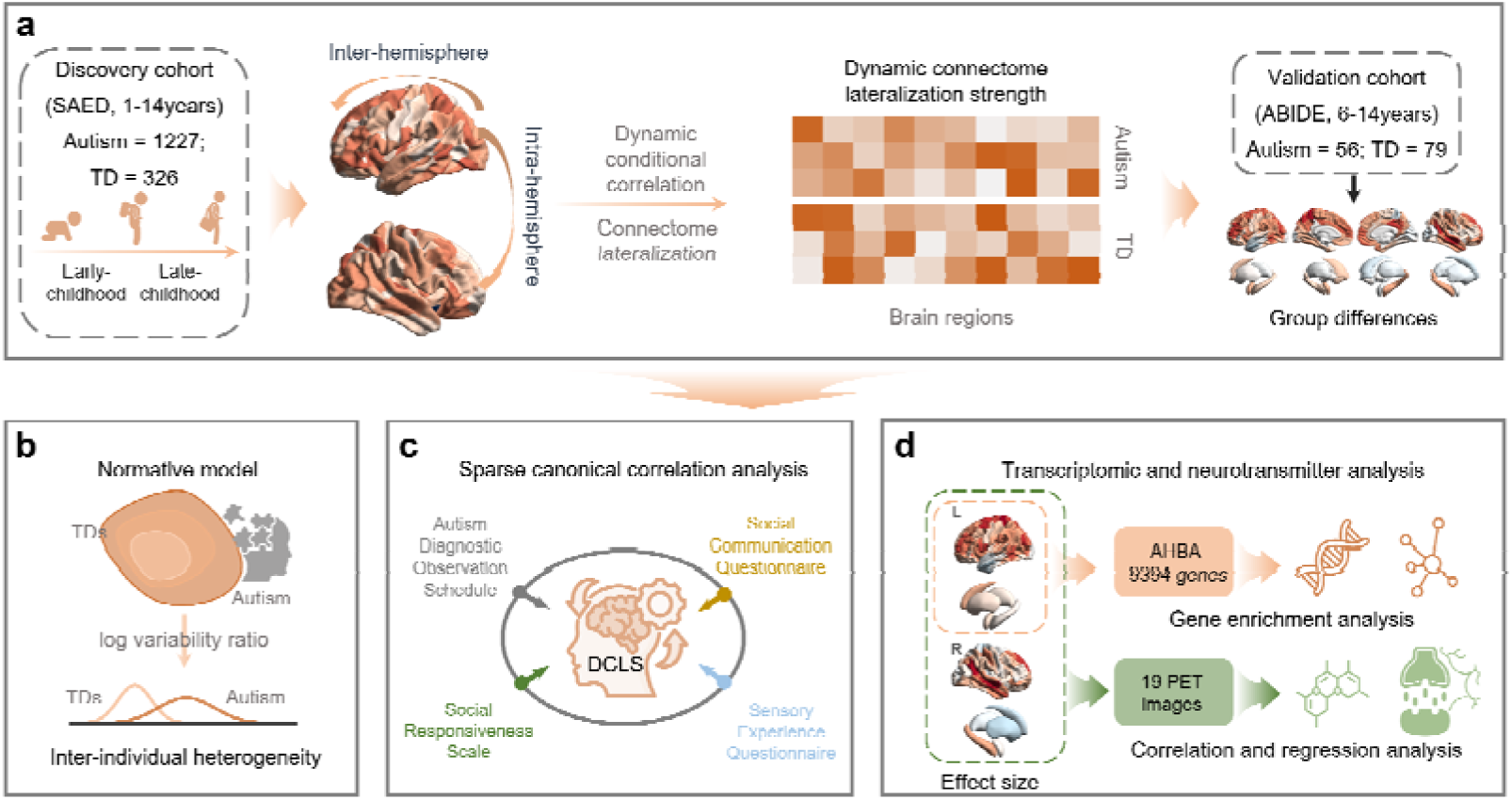
Analytical Framework. **a,** Estimating dynamic connectome lateralization strength (DCLS). Dynamic conditional correlation (DCC) was employed to construct dynamic connectomes; regional DCLS was quantified as the difference between inter- and intra-hemispheric functional connectivity strength. Spatiotemporal trajectories of DCLS were characterized across distinct developmental stages using the Shanghai Autism Early Developmental (SAED) cohort (discovery), with findings in late childhood validated using the Autism Brain Imaging Data Exchange (ABIDE) dataset (validation). **b,** Quantification of inter-individual heterogeneity. Normative modeling combined with the log variability ratio was utilized to assess difference in DCLS variability between autistic and TD individuals. **c,** Multivariate clinical associations. Sparse canonical correlation analysis (sCCA) was applied to delineate latent associations between whole-brain DCLS patterns and multidimensional clinical symptom profiles. **d,** Neurobiological decoding. The spatial map of DCLS alterations was spatially correlated with brain-wide gene expression signatures and neurotransmitter receptor/transporter distributions to elucidate underlying molecular substrates. autism, autistic individuals; TD, typically developing individuals.

### Atypical DCLS in different autistic developmental stages and hemispheres

The averaged DCLS maps across all autistic and TD individuals, as well as at each developmental stage, are shown in Supplementary Fig. 1. Compared with TD controls, autistic individuals exhibited significantly greater DCLS values (Cohen’s *d* > 0.2 and *p*_FDR_□<□0.05) in 69 brain regions across the entire childhood period (largest effect size: left precuneus = 0.412, Fig. 2a, Supplementary Table 3), 20 regions during early childhood (largest effect size: right superior temporal gyrus [STG] = 0.361, Fig. 2b, Supplementary Table 4), and 46 regions during late childhood (largest effect size: left precuneus = 0.476, Fig. 2c, Supplementary Table 5). Notably, no regions showed significantly higher DCLS in TD individuals than in autistic individuals. At the network level, during the entire period, autistic individuals exhibited higher DCLS values (mean Cohen’s *d* > 0.2) primarily in the left dorsal attention network (DAN), ventral attention network (VAN), and frontoparietal network (FPN), as well as in the right default mode network (DMN) (Fig. 2a). In the early childhood subgroup, increased DCLS was mainly observed in the left FPN and right DMN (Fig. 2b), whereas in the late childhood subgroup, higher DCLS was found in the left DAN, VAN, and FPN, as well as the bilateral DMN (Fig. 2c). Hemispheric comparison further revealed left-lateralized DCLS abnormalities in autism, which were significant during the entire period (*t*_(122)_ = 2.95, *p*_FDR_ □ = 0.006, Fig. 2a) and late childhood (*t*_(122)_ = 4.05, *p*_FDR_ □ = 4.4e-05, Fig. 2c), but not during early childhood (*t*_(122)_ = 1.85, *p*_FDR_ □ = 0.067, Fig. 2b). In the independent sample (ABIDE cohort), both the effect size brain map and the left-lateralized DCLS abnormalities observed in late childhood were successfully reproduced, showing a high spatial correlation (*r* = 0.421, *p* = 5.2e-12) and significant left-lateralization (*t*_(122)_ = 5.14, *p* = 1.0e-06) (Supplementary Fig. 2).

**Fig. 2.**
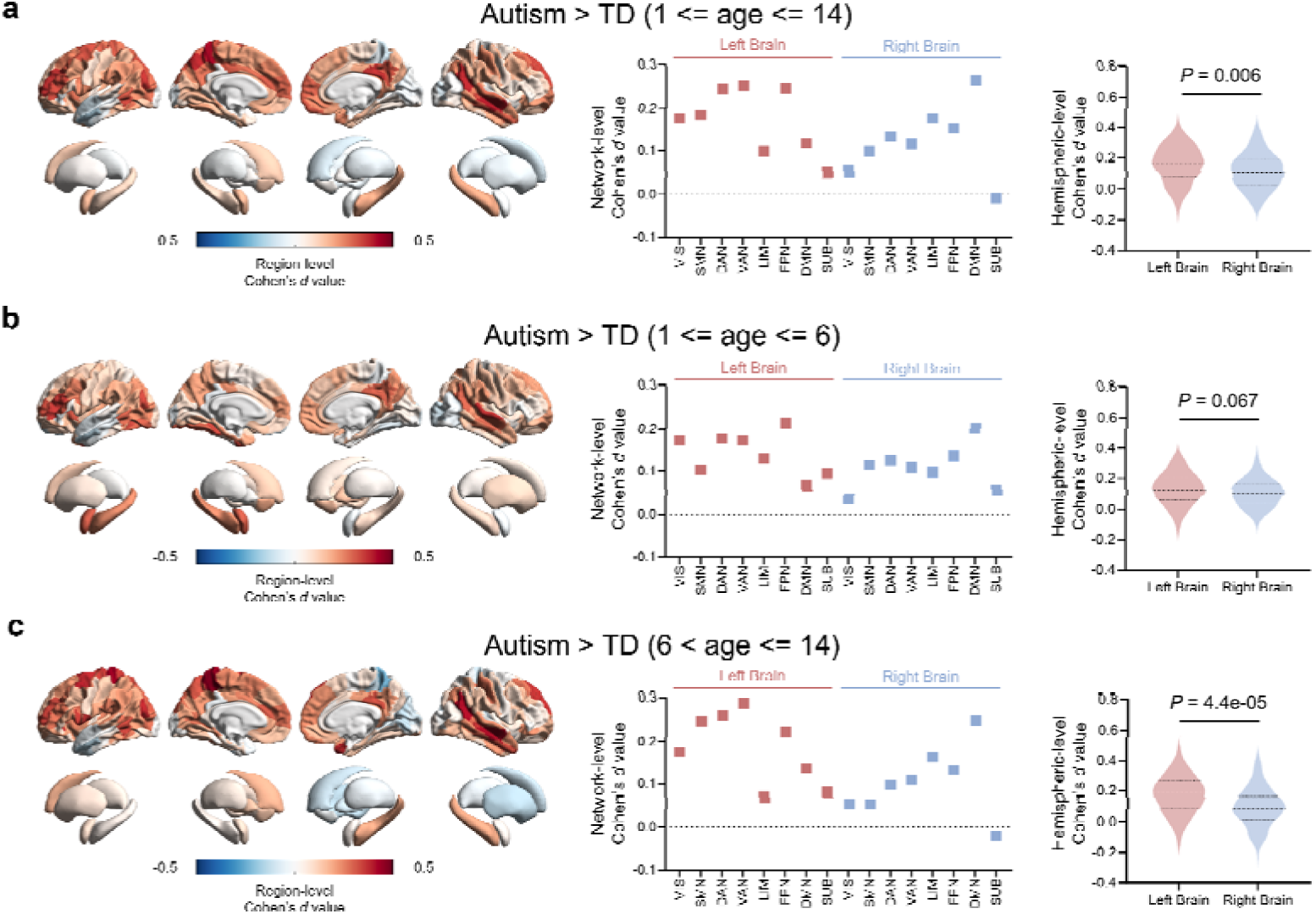
Group differences in dynamic connectome lateralization strength between autistic individuals and typically developing individuals. Effect sizes of group-level DCLS differences are displayed at the regional (left panels), network (middle panels), and hemispheric (right panels) levels during the entire childhood period (n_autism_ = 1227, n_TD_ = 326, **a**), early childhood (n_autism_ = 929, n_TD_ = 147, **b**), and late childhood (n_autism_ = 298, n_TD_ = 179, **c**). TD, typically developing individuals. **a-c,** The left panels show the regional effect sizes of group-level differences in DCLS are mapped onto a brain template. The middle panels show the average effect sizes of group-level differences on DCLS in each network. VIS, visual network; SMN, somatomotor network; DAN, dorsal attention network; VAN, ventral attention network; LIM, limbic network; FPN, frontoparietal network; DMN, default mode network; SUB, subcortical network. The right panels illustrate hemispheric comparisons of effect sizes of group-level differences in DCLS using two-sided paired-samples *t*-tests, and *p*-values were adjusted for multiple comparisons using the FDR correction. Data are presented as mean values (thick dashed lines) with quartiles (thin dashed lines).

### Heterogenous DCLS in different autistic developmental stages and hemispheres

Using individual-level DCLS deviations obtained from normative modeling, we further quantified the regional log variability ratio (lnVR) of these deviations between autistic and TD individuals across different developmental stages. Significant group difference was observed in only one brain region during early childhood (middle temporal gyrus, MTG, lnVR = 0.252, 95%CI = [0.004, 0.499], Fig. 3a), where autistic individuals showed greater DCLS variability than TD controls. In late childhood, this pattern became more widespread, with 14 regions showing higher variability in the autism group (largest lnVR: left parahippocampal gyrus = 0.414, 95%CI = [0.153, 0.680]), while 6 regions showing lower variability (smallest lnVR: right orbital gyrus = −0.337, 95%CI = [−0.600, −0.073]; Fig. 3b, Supplementary Table 6). At the network level, autistic individuals showed comparable DCLS variability to TD individuals during early childhood (|mean lnVR values| < 0.1, Fig. 3b), but exhibited greater variability during late childhood, particularly in the left hemisphere within the VAN (mean lnVR = 0.175) and LIM (mean lnVR = 0.183) (Fig. 3b). Hemispheric comparison in regional lnVR further revealed greater DCLS variability in the left hemisphere among autistic individuals, significant in both early childhood (*t*_(122)_ = 5.89, *p*_FDR_ = 3.4e-08, Fig. 3a) and late childhood (*t*_(122)_ = 12.65, *p*_FDR_ LJ< 1.0e-20, Fig. 3b). Moreover, the lnVR map showed a significant spatial correlation with the effect size map of case–control DCLS differences across the whole-brain during late childhood (*r* = 0.026, *p*_FDR_ = 0.686, Fig. 3d), but not during early childhood (*r* = 0.277, *p*_FDR_ = 2.0e-05, Fig. 3c). We further calculated the inter-subject similarity of whole-brain DCLS deviations among autistic individuals, which increased significantly with age during late childhood (*r* = 0.294, *p*_FDR_ = 4.9e-09, Fig. 3e), but not during early childhood (*r* = 0.054, *p*_FDR_ = 0.10).

**Fig. 3.**
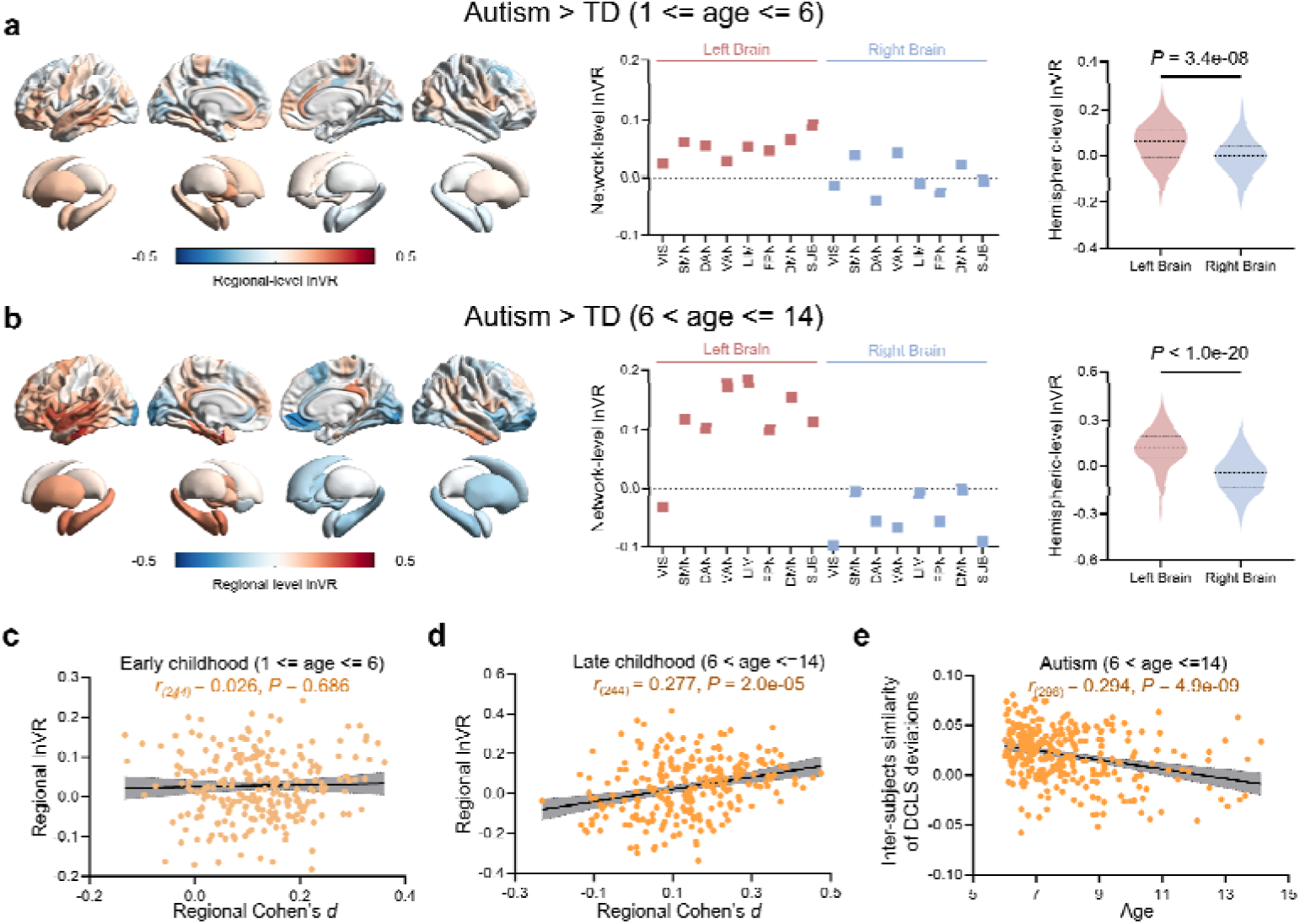
Log variability ratio (lnVR) of dynamic connectome lateralization strength deviations between autistic and typically developing individuals. The lnVR of DCLS deviations estimated from normative modeling are displayed at the regional (left panels), network (middle panels), and hemispheric (right panels) levels during early childhood (n_autism_ = 929, n_TD_ = 147, **a**), and late childhood (n_autism_ = 298, n_TD_ = 179, **b**). **a,b,** The left panels show the regional lnVR of DCLS deviations are mapped onto a brain template in early childhood (**a**) and late childhood (**b**). The middle panels show the average lnVR of DCLS deviations in each network. VIS, visual network; SMN, somatomotor network; DAN, dorsal attention network; VAN, ventral attention network; LIM, limbic network; FPN, frontoparietal network; DMN, default mode network; SUB, subcortical network. The right panels illustrate hemispheric comparisons of lnVR of DCLS deviations using two-sided paired-samples *t*-tests, and *p*-values were adjusted for multiple comparisons using the false-discovery-rate correction. Data are presented as mean values (thick dashed lines) with quartiles (thin dashed lines). **c,d,** The spatial association (Pearson’s correlations) between effect maps and lnVR map across whole brain regions during early childhood (**c**) and late childhood (**d**). *p*-values were adjusted for multiple comparisons using the FDR correction. **e,** The Pearson’s correlation between inter-subject similarity of whole-brain DCLS deviations among autistic individuals and their age during late childhood.

### Linked clinical measurements and DCLS in autistic individuals

To uncover the shared dimensions linking DCLS with clinical symptoms in the early and late childhood autism, we applied sCCA to relate regional DCLS features and clinical measurement profiles. Notably, there were no significant differences between early and late childhood autism groups on sex and Autism Diagnostic Observation Schedule (ADOS) total score. In the early childhood, the first sCCA model showed a significant correlation between DCLS features and clinical measurements (*r* = 0.748, *p*_perm-FDR_ < 0.001, Supplementary Fig. 3). The brain loading map revealed significant contributions from 20 regions (Fig. 4a and Supplementary Table 7), with the strongest regional loadings in the right amygdala (loading = 0.186, *p*_perm-FDR_ < 0.001), left superior temporal gyrus (loading = 0.181, *p*_perm-FDR_ < 0.001) and right orbital gyrus (loading = 0.151, *p*_perm-FDR_ = 0.006), and the highest network-level loading in the left limbic network (LIM, loading = 0.071, Fig. 4b). No significant hemispheric differences in brain loading strength were observed (*t*_(122)_ = −1.10, *p* = 0.274). This brain–clinical model was significantly associated with all the 17 clinical measurements (loadings > 0.100, *p*_perm-FDR_ < 0.01, Fig.4c), particularly Sensory Experience Questionnaire (SEQ) total scores (loading = 0.682, *p*_perm-FDR_ < 0.001), SEQ hypo-nonsocial scores (loading = 0.601, *p*_perm-FDR_ < 0.001), and SEQ hyper-social scores (loading = 0.584, *p*_perm-FDR_ < 0.001). In the late childhood, the first identified dimension remained significant (*r* = 0.970, *p*_perm-FDR_ < 0.001, Supplementary Fig. 3), with 32 regions contributing strongly (Fig. 4d and Supplementary Table 8), most prominently in the right amygdala (loading = 0.340, *p*_perm-FDR_ < 0.001), left middle frontal gyrus (loading = 0.316, *p*_perm-FDR_ < 0.001), left inferior parietal lobule (loading = 0.294, *p*_perm-FDR_ < 0.001) and left default mode network (DMN, loading = 0.166, Fig.4e). Moreover, a trend toward higher brain loadings in the left hemisphere was observed (*t*_(122)_ = 1.74, *p* = 0.043, one-sided paired-samples *t*-tests). The corresponding behavioral loading pattern indicated significant associations with 12 clinical measurements (Fig.4f), particularly Social Responsiveness Scale (SRS) mannerism scores (loading = 0.581, *p*_perm-FDR_ < 0.001), SRS total scores (loading = 0.558, *p*_perm-FDR_ < 0.001), and ADOS communication+social score (loading = 0.547, *p*_perm-FDR_ < 0.001).

**Fig. 4.**
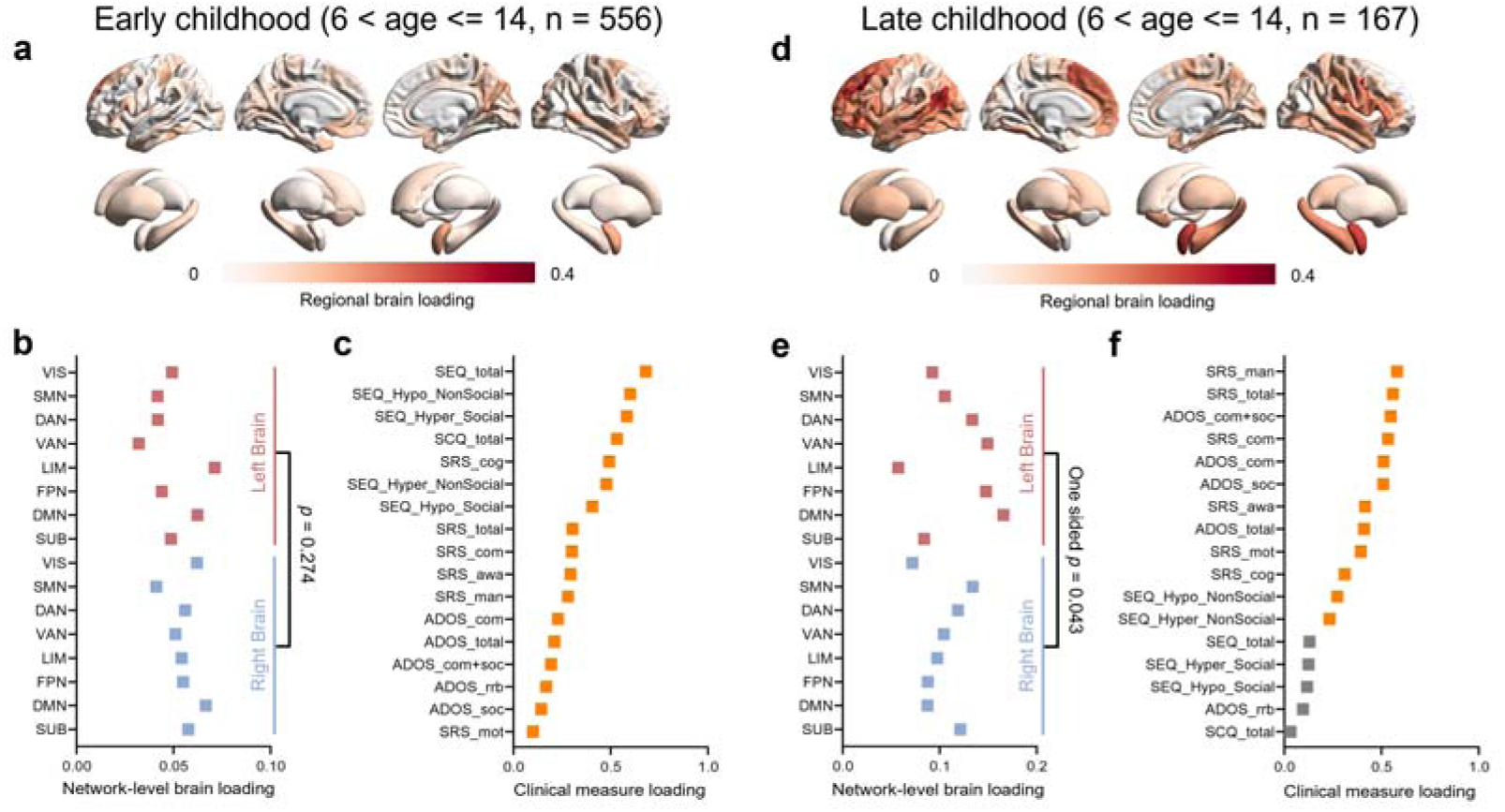
Brain–behavior associations between dynamic connectome lateralization strength and clinical measurements in autistic individuals. The results of sparse canonical correlation analysis (sCCA) in the early childhood (n_autism_ = 556, left panels, **a-c**) and late childhood (n_autism_ = 167, right panels, **d-f**). **a,d,** Regional brain loadings of the first canonical variate mapped onto a brain template. **b,e,** Average brain loadings of the first canonical variate in each network. The hemispheric comparisons were conducted for regional brain loadings using paired-samples *t*-test. VIS, visual network; SMN, somatomotor network; DAN, dorsal attention network; VAN, ventral attention network; LIM, limbic network; FPN, frontoparietal network; DMN, default mode network; SUB, subcortical network. **d,f,** Clinical loadings of the first canonical variate for each clinical measure. ADOS: Autism Diagnostic Observation Schedule; ADOS-com: ADOS communication; ADOS-soc: ADOS social interaction; ADOS-rrb: ADOS restricted and repetitive behaviors; ADOS-c+s: ADOS communication + social interaction; SCQ: Social Communication Questionnaire; SEQ: Sensory Experience Questionnaire; SEQ-hypoS: SEQ hypo-social; SEQ-hypoNS: SEQ hypo-nonsocial; SEQ-hypeS: SEQ hyper-social; SEQ-hypeNS: SEQ hyper-nonsocial; SRS: Social Responsiveness Scale; SRS-awa: SRS awareness; SRS-cog: SRS cognition; SRS-com: SRS communication; SRS-mot: SRS motivation; SRS-man: SRS mannerism. Yellow squares indicate significant associations (5,000 permutation tests followed by FDR correction), and gray squares indicate non-significant associations.

### Transcriptomic decoding of atypical DCLS across childhood subgroups in autism

By linking regional DCLS abnormalities to cortical gene expression profiles across childhood subgroups in autism, we aimed to identify the molecular substrates underlying atypical connectome lateralization. Specifically, partial least squares (PLS) regressions were first used to link the spatial patterns of DCLS group differences (i.e. regional Cohen’s *d* values) with 9,394 gene expressions from the AHBA for early and late childhood groups, respectively. A significant correlation was observed between the spatial pattern of the first PLS component (PLS1; Fig. 5a, explaining 22.3% of the variance in DCLS abnormalities; Fig. 5b) and the Cohen’s d map during late childhood (*r* = 0.472, *p*_spin_ = 0.003; Fig. 5c), whereas no significant association was found in early childhood (*r* = 0.368, *p*_spin_ = 0.086). Therefore, subsequent transcriptomic analyses focused on the late childhood stage. Using a one-sample Z test, genes were ranked according to their normalized PLS1 weights, yielding 2,385 significant genes (|Z| > 3, *p_FDR_*< 0.05, Fig. 5d) for enrichment analysis. These genes were primarily involved in biological processes related to the top 20 functional categories (Fig. 5e). Cell-type enrichment analysis indicated that the associated genes were enriched in multiple neuron types, including GABAergic, dopaminergic, glutamatergic and motor-related neurons (Fig. 5f). Moreover, human-disease-associated gene enrichment analysis showed that these genes are mainly enriched in memory impairment, visual seizure and cerebellar ataxia (Fig. 5g).

**Fig. 5.**
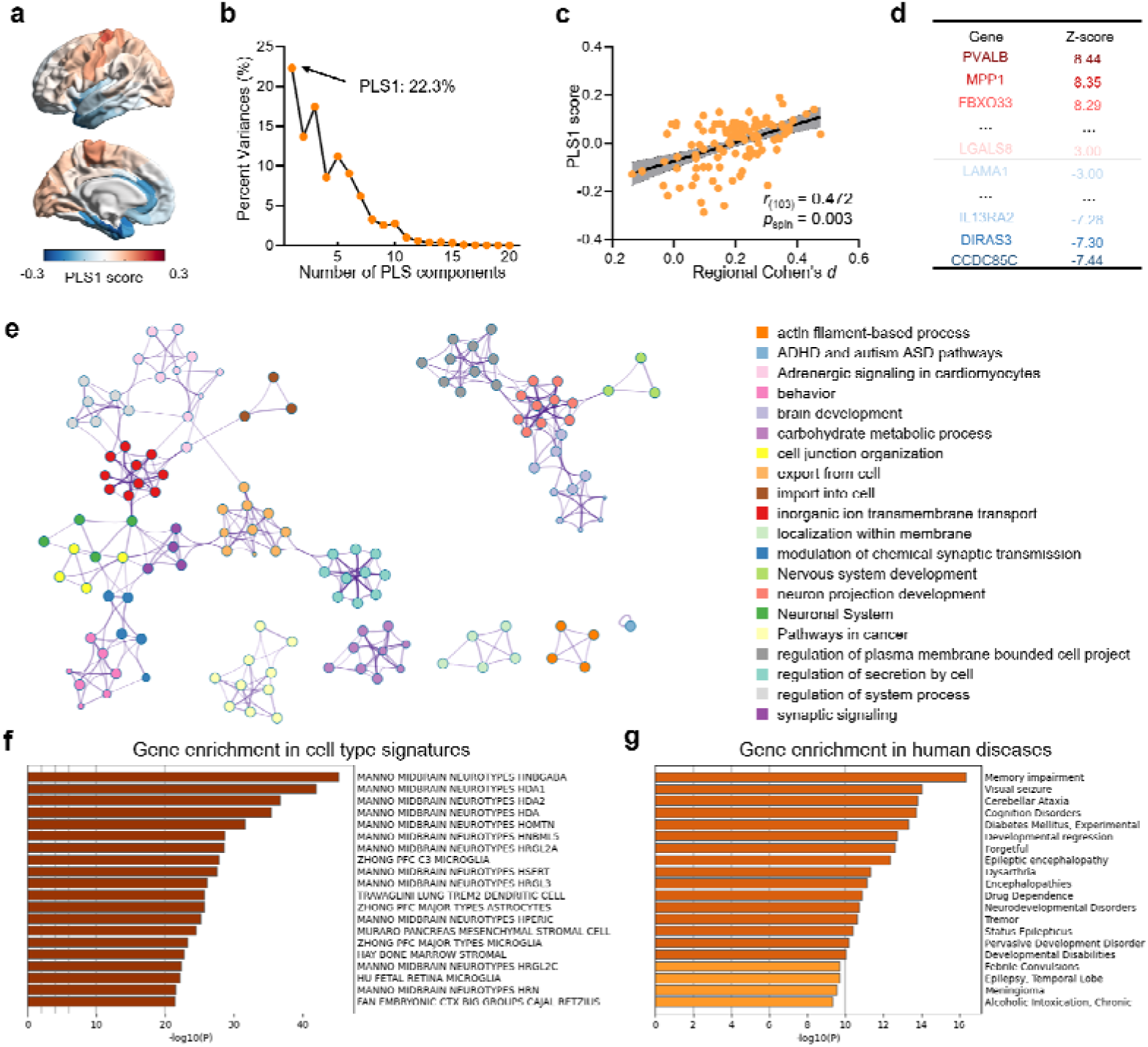
Partial least squares (PLS) correlation analysis between dynamic connectome lateralization strength abnormalities and human brain gene expression data during late childhood stage. **a,** Spatial distribution of the first partial least squares (PLS1) component in the left hemisphere. **b,** Variance explained by the first 20 PLS components derived from the regression analysis. **c,** Spatial correlation (Pearson’s correlation) between regional DCLS abnormalities (Cohen’s *d* value) and regional PLS1 scores. Statistical significance was evaluated using 5,000 times “spin”-based permutation test (*r*_(103)_ = 0.472, *p*_spin_ = 0.003, two-sided). **d,** Ranked PLS1 gene weights (|Z| > 3, *p*_FDR_ < 0.05). **e,** Top 20 gene ontology biological processes by enrichment analysis. Circle colors denote distinct functional categories, and circle sizes indicate the number of genes involved. **f,** Cell-type–specific enrichment of genes associated with DCLS abnormalities. **g,** Human disease–related gene enrichment associated with DCLS abnormalities. The Benjamini–Hochberg procedure was used for multiple comparison adjustments in **f** and **g**.

### Associations between neurotransmitter receptors and atypical DCLS in autism across developmental stages

To further elucidate the neurochemical basis of atypical connectome lateralization in autism across developmental stages, we correlated the spatial pattern of DCLS effect sizes (Cohen’s *d* values) with receptor density maps of 19 neurotransmitter systems across hemispheres and developmental stages. During early childhood, left-hemispheric DCLS abnormalities showed significant correlations with receptor densities of serotonin (5-HT_1A_ and 5-HTT), acetylcholine (α_4_β_2_ and M_1_), cannabinoid (CB_1_), norepinephrine (NET), and glutamate (mGluR_5_) receptors (*p*_spin-FDR_ < 0.05, Fig. 6a and Supplementary Table 9). A multiple linear regression (MLR) model incorporating these neurotransmitter systems significantly predicted the spatial distribution of left-hemispheric DCLS alterations (Fig. 6b, *R*^2^ = 0.130, *p* = 0.022), with 5-HT_1A_, NET, and α_4_β_2_ collectively accounting for over 75% of the explained variance (Fig. 6b). For the right hemisphere, DCLS abnormalities were significantly associated with serotonin (5-HT_2A_) and glutamate (mGluR_5_) receptors (*p*_spin-FDR_ < 0.05, Fig. 6a and Supplementary Table 9), and the corresponding MLR model significantly predicted the right-hemispheric DCLS pattern (Fig. 6c, *R*^2^ = 0.136, *p* = 1.5e-04) with 5-HT_2A_ contributing 54% to the model’s explanatory power (Fig. 6c). During late childhood, left-hemispheric DCLS abnormalities were significantly correlated with receptor densities of serotonin (5-HT_1B_, 5-HT_2A_ and 5-HTT), dopamine (D_1_ and D_2_), norepinephrine (NET), GABA_A_ and glutamate (NMDA and mGluR_5_) systems (*p*_spin-FDR_ < 0.05, Fig. 6a and Supplementary Table 10). The combined MLR model significantly predicted the spatial pattern of left-hemispheric DCLS alterations (Fig. 6d, *R*^2^=0.445, *p* = 2.9e-11), with 5-HT_2A_, 5-HTT, NET and mGluR_5_ showing the strongest relative contributions (Fig. 6d, explaining 77% of total variance). In the right hemisphere, DCLS abnormalities were significantly associated with serotonin (5-HT_1B_, 5-HT_2A_ and 5-HTT), dopamine (D_1_, D_2_ and DAT), norepinephrine (NET), GABA_A_, histamine (H_3_), cannabinoid (CB_1_), acetylcholine (α_4_β_2_ and VAChT) and glutamate (NMDA and mGluR_5_) systems (*p*_spin-FDR_ < 0.05, Fig. 6a and Supplementary Table 10). The corresponding MLR model achieved significant prediction performance (Fig. 6e, *R*² = 0.447, *p* = 5.9e-09), with 5-HT_2A_, with H_3_, GABA_A_, 5-HT_2A_, D_2_ and DAT showing the strongest relative contributions (Fig. 6e, explaining 74% of total variance).

**Fig. 6.**
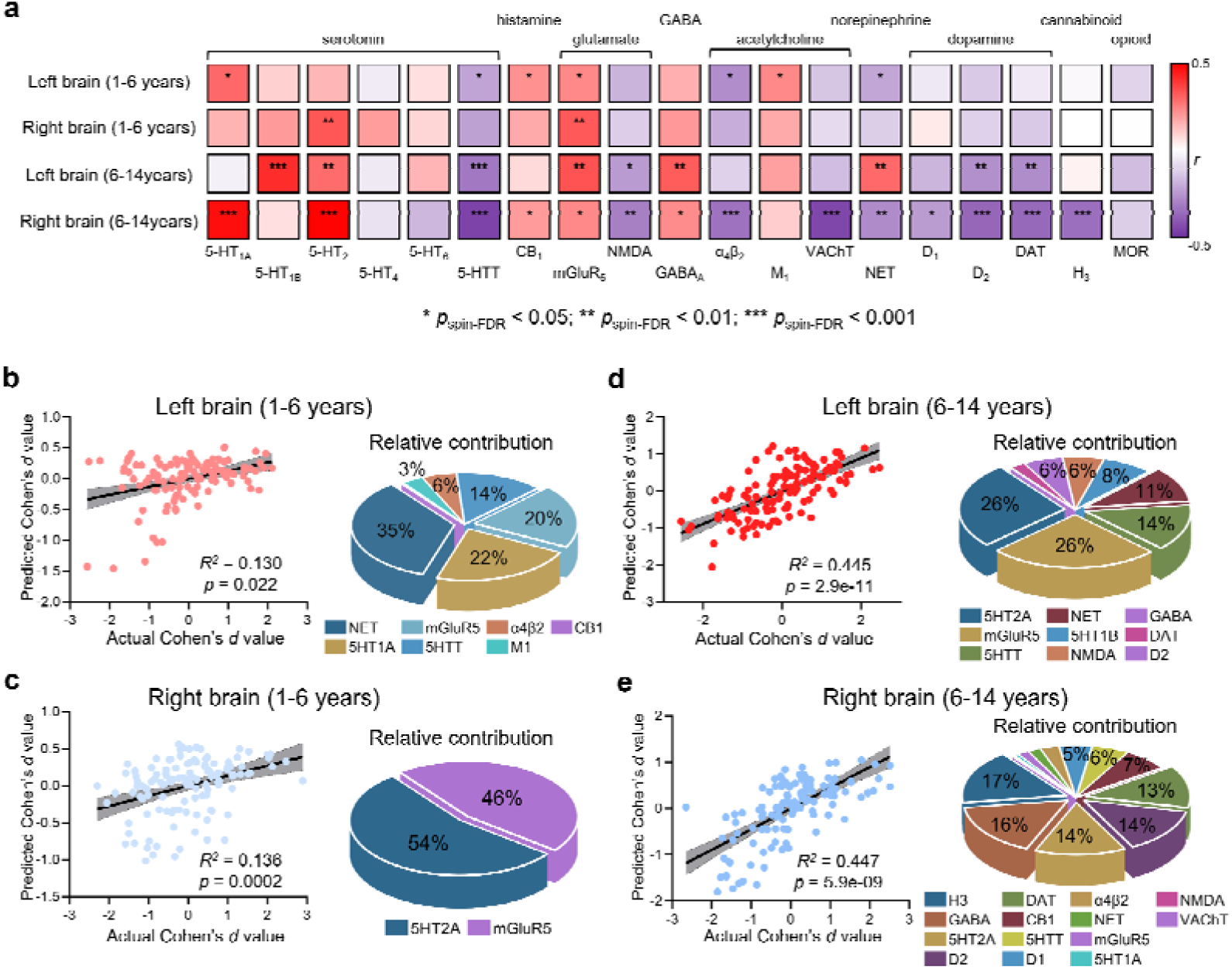
Associations between neurotransmitter receptor density and atypical DCLS patterns in autistic individuals. **a,** Spatial correlations (Pearson’s correlations) between DCLS effect size maps (Cohen’s d values) and receptor density maps across 19 neurotransmitter systems in the left and right hemispheres during early and late childhood. Statistical significance was assessed using 5,000 times “spin”-based permutation test (two-sided) followed by FDR correction. **b–e,** Multiple linear regression (MLR) models predicting the spatial distribution of hemispheric DCLS abnormalities from receptor density maps. The left panels show the prediction of DCLS abnormalities based on brain receptor distributions. The right panels show the relative contributions of each neurotransmitter system to the MLR model. **b,** Left hemisphere in early childhood (R² = 0.130, *p* = 0.022). **c,** Right hemisphere in early childhood (R² = 0.136, *p* = 1.5e-0.4). **d,** Left hemisphere in late childhood (R² = 0.445, *p* = 2.9e-11). **e,** Right hemisphere in late childhood (R² = 0.447, *p* = 5.9e-09).

## Discussion

This study addresses critical knowledge gaps regarding the developmental heterogeneity of connectome lateralization in autism across the full spectrum of childhood. Specifically, by leveraging an advanced dynamic connectome framework to contrast inter- and intra-hemispheric connectivity, we quantified individuals’ brain-wide dynamic connectome lateralization strength (DCLS) across two independent autism cohorts. Our findings provide converging evidence highlighting spatiotemporal, stage-dependent abnormalities and inter-individual heterogeneity on DCLS in children with autism that evolve distinctly from early to late childhood. Furthermore, we elucidated the multi-level neurobiological underpinnings of these DCLS abnormalities in autistic children by integrating clinical phenotypes, transcriptomic substrates, and neurochemical architecture. Overall, our discoveries offer a novel perspective on the spatiotemporal heterogeneity of connectome lateralization in autism across the entire span of childhood, with underlying implications for elucidating neurodevelopmental deviations and guiding stage-specific precision interventions at both macroscopic network and microscopic molecular levels.

Previous studies have documented abnormalities on static functional asymmetry^16^ and inter-hemispheric homotopic connectivity^18,45^ in autistic individuals older than 6 years. Extending these observations, our findings indicate a pervasive increase in hemispheric connectome lateralization across early and late childhood in autistic individuals, reflected by consistently elevated DCLS values relative to TD controls. The absence of regions showing reduced DCLS suggests a unidirectional alteration in connectome lateralization, pointing to excessive functional segregation between hemispheres in the autistic brain. This pattern converges with prior evidence of reduced inter-hemispheric communication^45^ and corpus callosum alterations^46,47^ in autism, and may reflect atypical developmental coordination between the two hemispheres during brain maturation. Importantly, our results further reveal a developmentally dynamic trajectory of connectome lateralization abnormalities in children with autism. In early childhood, increased DCLS in autism was spatially constrained, primarily involving the left FPN and the right DMN—networks that undergo rapid functional specialization during this developmental window^1,5^. As development progressed into late childhood, these abnormalities became increasingly widespread, extending across multiple large-scale networks, including the left FPN, DAN, VAN, as well as the bilateral DMN. This progressive expansion is consistent with a disruption in the typical maturation of inter-hemispheric integration, whereby early-emerging network-specific lateralization differences may be amplified and propagated across distributed systems as large-scale networks become more differentiated and hierarchically organized with age^48,49^, a process that appears to be independent of the severity of clinical symptoms. Notably, in late childhood, autistic individuals exhibited significantly greater DCLS abnormalities in the left hemisphere relative to the right, a pattern that was not observed during early childhood. This finding further indicates that connectome lateralization in autism is dynamically modulated across development rather than remaining static. Furthermore, the successful replication of these late-childhood, left-lateralized dysregulation patterns in autistic children within the independent ABIDE cohort attests to the robustness of these findings.

Beyond group-level mean differences, our analyses revealed abnormal inter-individual variability of DCLS in children with autism relative to TD controls, exhibiting a clear developmental progression that paralleled group-level effects. During early childhood, increased DCLS variability was highly circumscribed, limited to the left MTG. As a key hub for auditory and language processing, the left MTG has previously been shown to exhibit reduced activation during auditory language tasks in autistic individuals^50^, and reduced local activity has also been reported in preschool children with autism^51^, suggesting that the selective heterogeneity observed in this region may reflect early instability in the connectome lateralization of language-related circuitry. In contrast to this focal pattern, late childhood was characterized by a marked increase in both the spatial extent and magnitude of DCLS heterogeneity, spanning multiple cortical regions and large-scale networks including left VAN and LIM. Furthermore, across both early and late childhood, autistic individuals consistently exhibited greater heterogeneity in the left hemisphere, whereas the right hemisphere showed comparable or greater homogeneity relative to TD controls. Notably, whereas DCLS heterogeneity during early childhood was not associated with the magnitude of group-level connectome lateralization abnormalities, a significant correspondence emerged during late childhood, such that regions with larger mean deviations also exhibited greater inter-individual variability. Moreover, inter-individual heterogeneity among autistic children continued to increase with age during late childhood, indicating that connectome lateralization abnormalities become progressively more individualized over development. Together, these findings indicate that connectome lateralization abnormalities in autism encompass both shared developmental alterations and increasingly individualized neurobiological trajectories across childhood. Consistent with precision medicine frameworks that emphasize stratifying autism based on clinical and neurobiological profiles^28,52–55^, our results suggest that developmental stage– and individual-specific patterns of connectome lateralization may provide a useful basis for informing targeted and personalized intervention strategies.

Multivariate analyses further demonstrated that connectome lateralization abnormalities are meaningfully coupled with clinical phenotypes in children with autism, with the nature of these associations varying across development. In early childhood, brain–behavior coupling was dominated by limbic and temporal regions, including the amygdala and STG, and was most strongly related to sensory experience–related measures. This is consistent with evidence that atypical sensory hyper-/hypo-reactivity often emerges in infancy and toddlerhood (0–3 years, extending into 0–6 years), frequently preceding overt social deficits^56–58^. By late childhood, the brain–behavior dimension shifted toward a more distributed configuration involving left FPN, VAN, and DMN regions, networks that are central to social cognition, executive function, and integrative processing^59–61^. At this stage, lateralization abnormalities were more closely associated with core social–communicative symptoms, as indexed by SRS and ADOS measures. This transition corresponds to the increasing prominence of social-communication impairments and repetitive behaviors as children face greater demands for complex social navigation during school age^57^. Notably, in late childhood, these brain–behavior associations were accompanied by a trend toward left-hemispheric predominance in connectome lateralization. The observed left-hemispheric predominance may indicate that alterations in left-lateralized networks supporting language-related^62,63^ and integrative social-communicative processes^64^ become increasingly coupled to core symptom dimensions as development progresses^65^. These findings further underscore the developmental specificity of connectome lateralization–clinical phenotype coupling in autism and highlight the potential value of targeting lateralized network dynamics in developmental stage-appropriate interventions.

Furthermore, we elucidated the multi-level neurobiological underpinnings of DCLS abnormalities in children with autism by integrating transcriptomic substrates and neurochemical architecture. Our integrative analyses indicate that: (1) Transcriptomic decoding revealed significant associations between DCLS abnormalities and cortical gene expression profiles primarily in late childhood, where PLS1 explained a substantial portion of variance and highlighted genes enriched in transmembrane transport, chemical synaptic transmission and synaptic signaling. This aligns with the synaptic dysfunction hypothesis in autistic individuals, where aberrant synaptic pruning and connectivity maturation patterns during neurodevelopment contribute to core symptoms^66,67^, with late childhood marking a period of amplified molecular vulnerabilities as networks differentiate^68^. (2) Enriched genes were predominantly linked to GABAergic, dopaminergic, glutamatergic, and motoneuron cell types, consistent with evidence implicating neurotransmitter imbalances and neuronal dysregulation in autism etiology^69,70^. (3) Disease-associated enrichment pointed to memory impairment, visual seizures, and cerebellar ataxia, suggesting that developmental abnormalities in connectome lateralization are linked to molecular pathways supporting cognition^57^, sensory–motor integration^48,65^, and cerebellar development^71^ in autism. (4) Neurochemical analyses revealed stage- and hemisphere-specific associations between DCLS alterations and neurotransmitter systems that are highly relevant to autism. Early-childhood links with serotonergic, glutamatergic, and noradrenergic pathways align with their established roles in early cortical development^72^ and sensory regulation^73,74^ in autism. By late childhood, broader involvement of dopaminergic, GABAergic, and histaminergic systems may reflect emerging alterations in excitation–inhibition balance^75^, cognitive control, and behavioral regulation^76^, and underscore poly-neurotransmitter contributions to lateralized dysmaturation. Collectively, these multi-modal insights reveal the evolving molecular and neurochemical foundations of connectome lateralization abnormalities in autism, from early constrained to late widespread patterns. This points toward developmentally informed strategies that target these substrates to address heterogeneous clinical trajectories, while also highlighting the need for future research to consider the influence of comorbidity and genetic overlap at different developmental stages^77^.

Despite the strengths of our large sample size, developmental perspective and multi-modal analyze pipeline, several limitations should be noted. First, the cross-sectional design limits inferences about intra-individual developmental trajectories; longitudinal studies are needed to confirm the progression of connectome lateralization abnormalities and their links to clinical phenotypes. Second, we related the connectome lateralization abnormalities map in autistic children with the gene expression and neurochemical brain map sourced from healthy adults. The observed associations may not fully capture autism-specific or developmentally dynamic molecular and neurochemical alterations, and therefore warrant validation using disorder- and age-matched datasets. Third, our sample was restricted to early and late childhood (1–14 years), limiting the generalizability of the findings to adolescence and adulthood, during which connectome lateralization and clinical phenotypes may undergo further reorganization. Future studies spanning the full lifespan will be essential to determine whether the developmental patterns observed here persist, stabilize, or diverge at later stages.

In summary, our study provides a comprehensive characterization of the spatiotemporal and inter-individual heterogeneity of connectome lateralization in autism across childhood. We demonstrate that lateralization abnormalities evolve from early, focal disruptions to widespread, individualized patterns by late childhood, and that these alterations are meaningfully associated with clinical phenotypes, transcriptomic profiles, and neurotransmitter systems. These findings highlight the developmental specificity of connectome lateralization and underscore its potential as a biomarker for developmental stage- and individual-specific intervention strategies. Together, our results offer novel insights into the neurodevelopmental mechanisms underlying autism in the first 14 years and provide a framework for leveraging dynamic connectome features in precision neuroscience.

## Methods

### Discovery dataset - SAED cohort

The discovery dataset enrolled 1299 autistic individuals (range: 1-14 years) and 349 TD individuals (range: 1-14 years) from the SAED Cohort^15,38^. We then applied the following exclusion criteria for quality control: (1) missing information on sex or age; (2) missing data of sMRI scan or rsfMRI scan; (3) excessive head motion (mean FD > 0.5). After quality control, 1227 autism and 326 TD individuals were divided into two developmental groups: early childhood (1 ≤ age ≤ 6 years) and late childhood (6 < age ≤ 14 years). The early childhood group consisted of 929 autistic individuals (768 males, mean age ± SD = 3.67 ± 1.13 years) and 147 TD individuals (68 males, mean age ± SD = 3.94 ± 1.37 years). The late childhood group consisted of 298 autistic individuals (245 males, mean age ± SD = 7.93 ± 1.68 years) and 179 TD individuals (70 males, mean age ± SD = 8.72 ± 1.79 years). All clinical diagnoses of autism were conducted by professional pediatricians based on Diagnostic and Statistical Mental Disorders, Fifth Edition (DSM-5) and ADOS or ADOS-Toddler module. Additionally, clinical measurements related to autism were administered by either pediatricians or their guardians, including the ADOS, SCQ, SEQ, and SRS. The total scores and subitem scores were used for subsequent analyses. Detailed demographic characteristics and assessments information are provided in Supplementary Table 1. This study was approved by the Ethics Committee of Xinhua Hospital affiliated with the Shanghai Jiao Tong University School of Medicine (XHEC-C-2019-076), and all participants provided written informed consent by their guardians.

### External validation data - ABIDE cohort

An independent cohort of late childhood participants (6 < age ≤ 14 years; 56 autism and 97 TD) was selected from six sites of ABIDE-I and II^30,31^, according to the following inclusion criteria: (1) complete information on sex and age; (2) complete data of sMRI scan and rsfMRI scan; (3) head motion (mean FD) < 0.5; (4) site sample size > 10 individuals; (5) rsfMRI scan duration > 6 minutes. The demographic characteristics of all included sites and participants are presented in Supplementary Table 2.

### MRI data acquisition and preprocessing

Participants in the SAED cohort underwent at least a T1-weighted structural scan and a rsfMRI scan on one of two MRI scanners: a Siemens Verio 3.0T MRI scanner (849 autism and 278 TD individuals) or a Philips Ingenia 3.0T MRI scanner (378 autism and 48 TD individuals), which was served as site information for subsequent analyses. Detailed scan parameters of SAED and ABIDE cohort are provided in Supplementary Table 11 and Supplementary Table 12. All sMRI and rsMRI data from the two cohorts were preprocessed using a standardized fMRIPrep (v.20.2.1) protocol^78^ (see Supplementary Methods), followed by a spatial smoothing (6 mm full-width at half-maximum) and 0.01-0.08 Hz band-pass filtering for the rsfMRI data using Data Processing & Analysis of Brain Imaging v.9.0 (DPABI v.9.0)^79^.

### Estimation of dynamic connectome lateralization strength

Fig. 1a illustrates the pipeline employed for the DCLS analysis. Firstly, we extracted the average blood-oxygen-level-dependent (BOLD) signals for 210 cortical and 36 subcortical regions defined in Human Brainnetome Atlas^80^, with each hemisphere containing 123 regions. These regions were categorized into eight brain networks per hemisphere: VIS, SMN, DAN, VAN, LIM, FPN, DMN and SUB. Thereafter, dynamic conditional correlation (DCC) was applied to calculate the dynamic connectome matrixes for each time point^28,39,40^, resulting in the generation of a participant-specific connectome as,

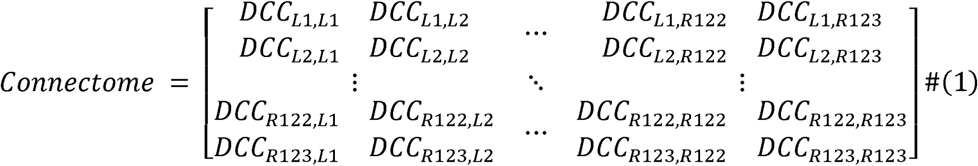

where *DCC_Li,Rj_* is Fisher-z-transformed DCC coefficient between the BOLD signals of left region *i* and right region *j*. Taking the first regions in the left and right hemisphere as examples, the DCL values at time *t* is defined as,

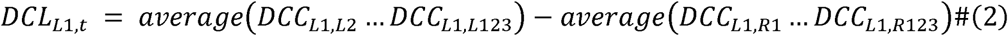

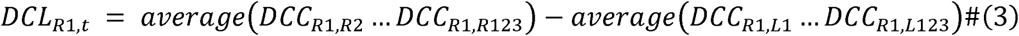

In this study, only correlations greater than 0.2 were considered to filter out weak or noise-related connectivity, excluding negative correlations^81,82^. In addition, focusing on the strength of individual brain functional lateralization and aiming to minimize the influence of different scanning time points on functional connectivity estimates^26^, we further calculated the time-averaged DCL (i.e. dynamic connectome lateralization strength, DCLS) for each region for subsequent analysis. Thus, a positive DCLS value signifies that this region exhibits stronger functional synchronizations with other regions in the ipsilateral hemisphere than the functional synchronizations with all regions in the contralateral hemisphere. To ensure comparability of DCLS values across imaging scanners or sites, ComBat harmonization was applied to both the SAED and ABIDE cohorts, with sex, age, and mean framewise displacement included as covariates in the models^83^. A two-sample t-test was performed to compare the mean difference in regional DCLS between autism and TD individuals, and regional Cohen’s *d* values were further estimated. False discovery rate correction was conducted for a total of 246 regional-level multiple comparisons. Furthermore, the cumulative effect size (quantified by Cohen’s *d*) within each hemispheric network was calculated, and the interhemispheric difference was conducted by paired t-test.

### Heterogeneity measures in region-level and whole brain-level DCLS

To investigate whether individuals with autism exhibit greater heterogeneity than control group in DCLS, we first established a childhood-stage-specific normative model to quantify the DCLS deviations of each autistic individual relative to the TD group of corresponding childhood stage^32^, and then used the log variability ratio (lnVR) to evaluate the relative heterogeneity between the autism and TD groups^41^ (Fig. 1b). Specifically, the DCLS deviation of region *i* for an autistic individual *j* was defined as,

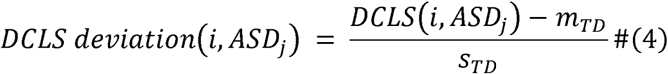

where *DCLS(i, autism_j_)* denotes the DCLS value of region *i* for individual *j*, *m*_TD_ and *s*_TD_ are the mean and s.d. DCLS value of the same-age TD group. For TD individuals, the DCLS deviation was defined as,

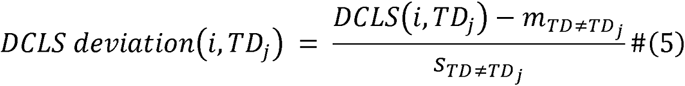

where *DCLS(i, TD_j_)* is the DCLS value of region *i* for individual *j*, *m*_TD≠TDj_ is the mean DCLS value of the same-age TD group excluding individual *j*, and s_TD≠TDj_ is the s.d. of the same-age TD group except individual *j*. Based on the DCLS deviation value for each individual, the regional lnVR of DCLS is given by the following equation,

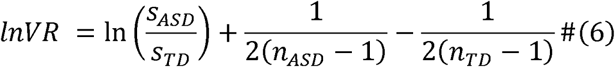

where *s*_autism_ and *s*_TD_ are the group s.d. of DCLS, and *n*_autism_ and *n*_TD_ are the group sizes for autism and TD groups, respectively. The positive lnVR signifies that autistic individual exhibit greater DCLS heterogeneity than TD group in this region. Statistical significance of lnVR was determined based on its sampling variance estimated via the Delta method (trigamma approximation)^84–86^. Similarly, the cumulative lnVR within each hemispheric network was calculated, and the interhemispheric difference was conducted by paired *t*-test. Additionally, to examine whether whole-brain heterogeneity in DCLS deviation among autistic individuals increases with age, we computed the mean Pearson’s correlation coefficients of the 246 regional DCLS deviation values between each autistic individual and all other autism participants within the early-childhood and late-childhood groups, respectively. We then evaluated the association between inter-individual similarity of DCLS deviations and age.

### Sparse canonical correlation analysis between DCLS and clinical measurements

We performed a sparse canonical correlation analysis (sCCA)^42–44^ to delineate latent dimensions linking functional lateralization features with clinical measurements in early and late childhood autism groups, separately. Unlike traditional canonical correlation analysis, sCCA imposes sparsity constraints on both neuroimaging and behavioral data matrices, which enhances model interpretability, improves feature selection specificity, and mitigates overfitting in high-dimensional settings. Specifically, we utilized sCCA to identify coupled patterns by maximizing the canonical correlations between functional lateralization features and clinical measurements, thereby uncovering symptom dimensions most strongly aligned with underlying neurobiological variability. The optimization objective is defined as,

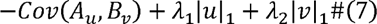

where *A* and *B* are the data matrices for regional DCLS features and clinical measurements, *u* and *v* denote their associated loading vectors, and λ_1_ and λ_2_ are the sparsity regularization terms applied to the loadings. We employed a grid search procedure in the range of [0.1, 0.2, 0.3, 0.4, 0.5, 0.6, 0.7, 0.8, 0.9, 1] to determine the optimal sparsity parameters for computing the canonical correlations (Supplementary Fig. 3). Next, we computed Pearson’s correlations between each original variable and its corresponding canonical variate to get brain and clinical feature loadings. The statistical significance of the resulting canonical correlations and feature loadings were evaluated using 5,000 permutation tests followed by FDR correction. Finally, to facilitate interpretation of each significant canonical variate, the absolute loading values of DCLS features and clinical measurements were reported, with higher loadings indicating a greater contribution of individual features to the corresponding canonical variates^42^.

### Imaging transcriptome analysis

We extracted brain-wide gene expressions of six postmortem human brains from the AHBA transcriptomic dataset^35,36^ (http://human.brain-map.org). Since only two donors have available data in the right hemisphere, and further excluding genes with low level of similarity across donors (< 0.2), resulting in a final cortical expression level matrix of 105 regions × 9394 genes. Then, partial least squares (PLS) regression was conducted to investigate the association between transcriptome and regional DCLS abnormalities (i.e., regional Cohen’s *d* values of DCLS comparisons between autism and TD) in early and late childhood autism groups, respectively. Gene expression levels were treated as predictor variables to predict regional DCLS abnormalities across 105 cortical regions in the left hemisphere. The first latent component (PLS1) captured the cortical expression pattern most strongly associated with the spatial distribution of DCLS alterations. To account for spatial autocorrelation, a 5,000 times *“spin”*-based permutation test was applied^87^. The contribution of each gene to PLS1 was further quantified using a bootstrapping approach. For each gene, a Z-score was calculated by dividing its weight by the corresponding bootstrap-derived standard error, and all 9,394 genes were subsequently ranked based on these Z-scores. Genes with |Z| > 3 and a *p*_FDR_ value below 0.05 were retained for gene enrichment analysis by Metascape (v.3.5)^88^.

### Neurotransmitter receptors and transporters analysis

We further investigated the spatial correlations between regional DCLS abnormalities and 9 distinct neurotransmitter systems, including dopamine (D_1_, D_2_ and DAT), norepinephrine (NET), serotonin (5-HT_1A_, 5-HT_1B_, 5-HT_2_, 5-HT_4_, 5-HT_6_ and 5-HTT), acetylcholine (α4β2, M_1_,VAChT), glutamate (mGluR_5_ and NMDA), γ-aminobutyric acid (GABA_A_), histamine (H_3_), cannabinoid (CB_1_) and opioid (MOR) (detailed information see Supplementary Table 13), which derived from previously reported PET images^37^. Specifically, we first averaged the PET images of each neurotransmitter receptor and transporter across participants and extracted the mean neurotransmitter expression values for 246 brain regions defined by the Human Brainnetome Atlas. We then assessed the spatial correlation (Pearson’s correlation) between regional DCLS abnormalities and each neurotransmitter receptor/transporter in the left and right hemispheres and for the early- and late-childhood groups separately. Statistical significance was evaluated using 5,000 times *“spin”*-based permutation tests followed by FDR correction^87^. Subsequently, the densities of neurotransmitter receptors and transporters that showed significant associations were used to predict regional DCLS abnormalities using a multiple linear regression model, performed separately for the left and right hemispheres and for the early- and late-childhood groups. Additionally, the relative contribution of each neurotransmitter system to the multiple linear regression model was quantified by calculating the proportion of the squared standardized coefficient to the total sum of squared coefficients across all predictors^89^.

## Acknowledgements

This study was supported by grants from the National Natural Science Foundation of China (82430104, 82125032), the China Brain Initiative Grant (STI2030-Major Projects 2021ZD0200800), the Science and Technology Commission of Shanghai Municipality (YDZX20253100003001, 23Y21900500, 23DZ2291100 and 2018SHZDZX01), the Shanghai Municipal Commission of Health and Family Planning (GWVI-11.1-34, 2020CXJQ01, 2018YJRC03), Innovative research team of high-level local universities in Shanghai (SHSMU-ZDCX20211100), “Discipline Peak-Climbing Plan” of Xinhua Hospital Affiliated to Shanghai Jiao Tong University School of Medicine (XKPF2024A50011), Humanity and Social Science Foundation of the Ministry of Education of China (24YJC190046), and Special Fund for Basic Scientific Research of Central Colleges (ZYGX2021J036).

## Authorship Contribution Statement

Q.L., Q.Y.L., W.Z. and F.L. conceptualized and designed the project. Q.L., Q.Y.L., W.Z. and F.L. drafted the original paper. Q.L., Q.Y.L., X.L., X.W. and X.Z. performed the statistical analysis. W.Z., L.Z., T.R., C.H., H.T, L.H., K.L., J.C., W.X. and Q.Z. contributed to the acquisition of research data. K.M.K. critically reviewed the paper for important intellectual content. W.Z. and F.L. supervised the project. All author reviewed the final manuscript.

## Data Availability

The MRI data and phenotypic information of discovery cohort (SAED cohort) that support the findings of this study are available on request from the corresponding authors. The MRI data of validation cohort (ABIDE I and II) are publicly available at http://fcon_1000.projects.nitrc.org/indi/abide/abide_I.html and https://fcon_1000.projects.nitrc.org/indi/abide/abide_II.html. The transcriptomic dataset of 6 postmortem adult brains can be found in the Allen Brain Atlas (https://human.brain-map.org/static/download). Gene enrichment analysis was performed using Metascape (v.3.5) (https://metascape.org). Neurotransmitter maps are publicly available at https://github.com/netneurolab/hansen_receptors.

## Code Availability

Brain images were processed by using fMRIPrep (v.20.2.1) (https://fmriprep.org) and DPABI (v.9.0) (http://rfmri.net/dpabi). Gene enrichment analysis was performed using Metascape (v.3.5) (https://metascape.org). The visualization of brain mapping images was conducted using the ENIGMA Toolbox (v.2.0.0) (https://enigma-toolbox.readthedocs.io/en/latest/). The codes used in this study are available on request from the corresponding authors.

