## Supplementary Figure 1-3 and Supplementary Table 1-13 for "Connectome lateralization in autism across the first 14 years: heterogeneity related to developmental stage, hemisphere, and pathophysiology"

Number of Supplementary Figures: 3
Number of Supplementary Tables: 13

**MRI Data Preprocessing**

All structural MRI and resting-state fMRI (rsfMRI) data from two datasets (ABIDE cohort & UESTC cohort) were preprocessed using fMRIPrep 20.2.1[1,2], which is based on Nipype 1.5.1[3,4].

The T1-weighted (T1w) image was corrected for intensity non-uniformity (INU) with N4BiasFieldCorrection, distributed with ANTs 2.3.3, and used as T1w-reference throughout the workflow. The T1w-reference was then skull-stripped with a Nipype implementation of the antsBrainExtraction.sh workflow (from ANTs), using OASIS30ANTs as target template. Brain tissue segmentation of cerebrospinal fluid (CSF), white-matter (WM) and gray-matter (GM) was performed on the brain-extracted T1w using fast. Volume-based spatial normalization to one standard space (MNI152NLin2009cAsym) was performed through nonlinear registration with antsRegistration (ANTs 2.3.3), using brain-extracted versions of both T1w reference and the T1w template. The following template was selected for spatial normalization: ICBM 152 Nonlinear Asymmetrical template version 2009c.

For the resting-state fMRI data, the first 5 volumes were discarded to ensure steady-state longitudinal magnetization. Then, a reference volume and its skull-stripped version were firstly generated using a custom methodology of fMRIPrep. Susceptibility distortion correction (SDC) was omitted. The BOLD reference was then co-registered to the T1w reference using flirt with the boundary-based registration cost-function. Co-registration was configured with nine degrees of freedom to account for distortions remaining in the BOLD reference. Head-motion parameters with respect to the BOLD reference (transformation matrices, and six corresponding rotation and translation parameters) are estimated before any spatiotemporal filtering using mcflirt. BOLD runs were slice-time corrected using 3dTshift from AFNI 20160207. The BOLD time-series (including slice-timing correction when applied) were resampled onto their original, native space by applying the transforms to correct for head-motion. These resampled BOLD time-series will be referred to as preprocessed BOLD in original space, or just preprocessed BOLD. The BOLD time-series were resampled into standard space, generating a preprocessed BOLD run in MNI152NLin2009cAsym space. First, a reference volume and its skull-stripped version were generated using a custom methodology of fMRIPrep. Several confounding time-series were calculated based on the preprocessed BOLD: framewise displacement (FD), DVARS and three region-wise global signals. FD was computed using two formulations following Power (absolute sum of relative motions) and Jenkinson (relative root mean square displacement between affines). FD and DVARS are calculated for each functional run, both using their implementations in Nipype. The three global signals are extracted within the CSF, the WM, and the whole-brain masks. Additionally, a set of physiological regressors were extracted to allow for component-based noise correction. Principal components are estimated after high-pass filtering the preprocessed BOLD time-series (using a discrete cosine filter with 128s cut-off) for the two CompCor variants: temporal (tCompCor) and anatomical (aCompCor). tCompCor components are then calculated from the top 2% variable voxels within the brain mask. For aCompCor, three probabilistic masks (CSF, WM and combined CSF+WM) are generated in anatomical space. The implementation differs from that of Behzadi et al. in that instead of eroding the masks by 2 pixels on BOLD space, the aCompCor masks are subtracted a mask of pixels that likely contain a volume fraction of GM. This mask is obtained by thresholding the corresponding partial volume map at 0.05, and it ensures components are not extracted from voxels containing a minimal fraction of GM. Finally, these masks are resampled into BOLD space and binarized by thresholding at 0.99 (as in the original implementation). Components are also calculated separately within the WM and CSF masks. For each CompCor decomposition, the k components with the largest singular values are retained, such that the retained components' time series are sufficient to explain 50 percent of variance across the nuisance mask (CSF, WM, combined, or temporal). The remaining components are dropped from consideration. The head-motion estimates calculated in the correction step were also placed within the corresponding confounds file. The confound time series derived from head motion estimates and global signals were expanded with the inclusion of temporal derivatives and quadratic terms for each. Frames that exceeded a threshold of 0.5 mm FD or 1.5 standardized DVARS were annotated as motion outliers. All resampling can be performed with a single interpolation step by composing all the pertinent transformations (i.e. head-motion transform matrices, susceptibility distortion correction when available, and co-registrations to anatomical and output spaces). Gridded (volumetric) resampling was performed using antsApplyTransforms (ANTs), configured with Lanczos interpolation to minimize the smoothing effects of other kernels. Non-gridded (surface) resampling was performed using mri_vol2surf (FreeSurfer).

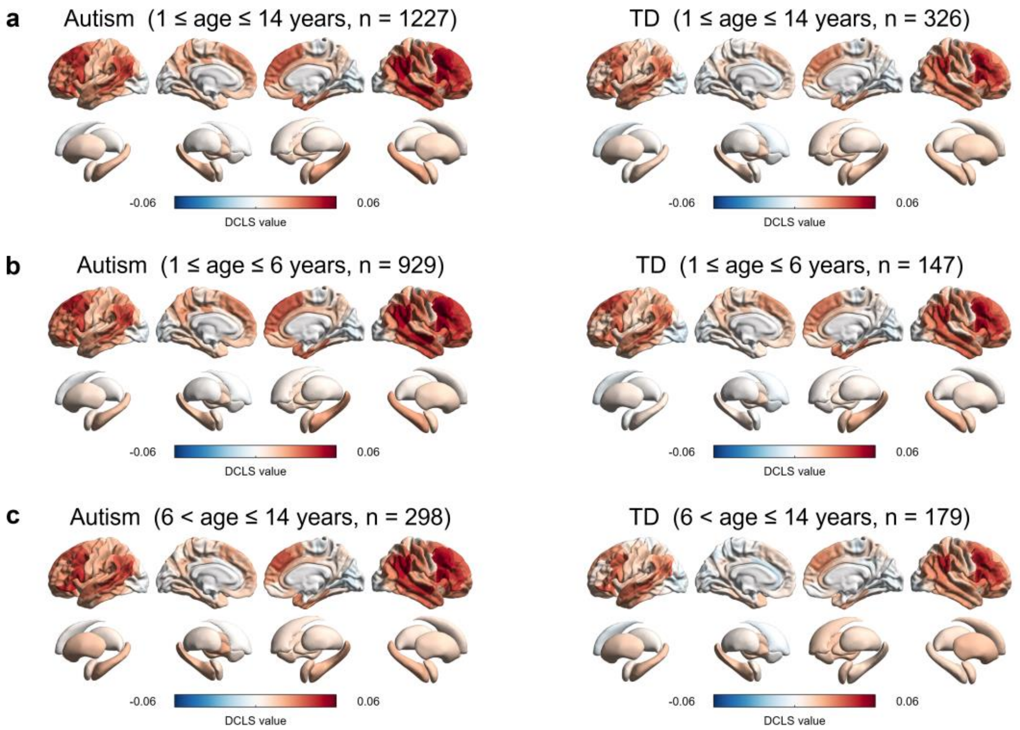


**Supplementary Fig. 1. The averaged dynamic connectome lateralization strength (DCLS) maps across all individuals in each group of SAED cohort. a** DCLS map across all autism and TD individuals (0 < age ≤ 14 years). **b** DCLS map across all autism and TD individuals in early childhood stage (0 < age ≤ 6 years). **c** DCLS map across all autism and TD individuals in late childhood stage (6 < age ≤ 14 years). A positive DCLS value signifies that this region exhibits stronger functional synchronization with other regions in the ipsilateral hemisphere than the functional synchronization with all regions in the contralateral hemisphere. TD: typically developing individuals; DCLS: dynamic connectome lateralization strength.

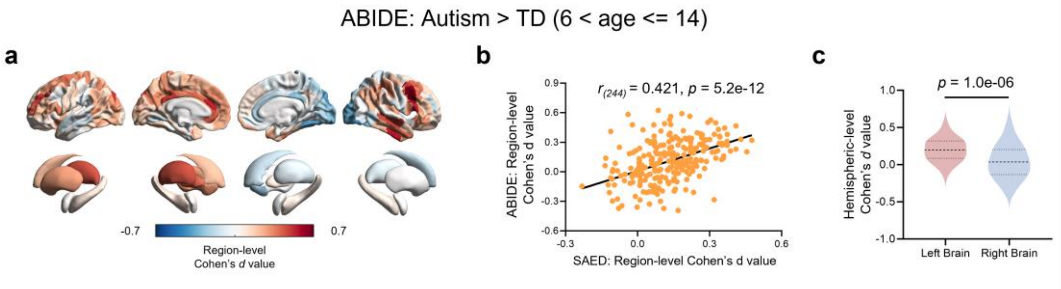


**Supplementary Fig. 2. Replication of dynamic connectome lateralization strength (DCLS) abnormalities with the independent ABIDE cohort ($n_{autism}$=56, $n_{TD}$=97). a,** Regional effect sizes of group-level differences in DCLS are mapped onto a brain template. **b,** The Person's correlation between the regional effect sizes in discovery data (SAED) and validation (ABIDE) datasets. SAED: Shanghai Autism Early Developmental Cohort**;** ABIDE: Autism Brain Imaging Data Exchange Cohort. **c,** Comparison of effect sizes between the left and right hemispheres using a two-sided paired-samples t-test. Data are shown as mean values (thick dashed line) with quartiles (thin dashed lines).

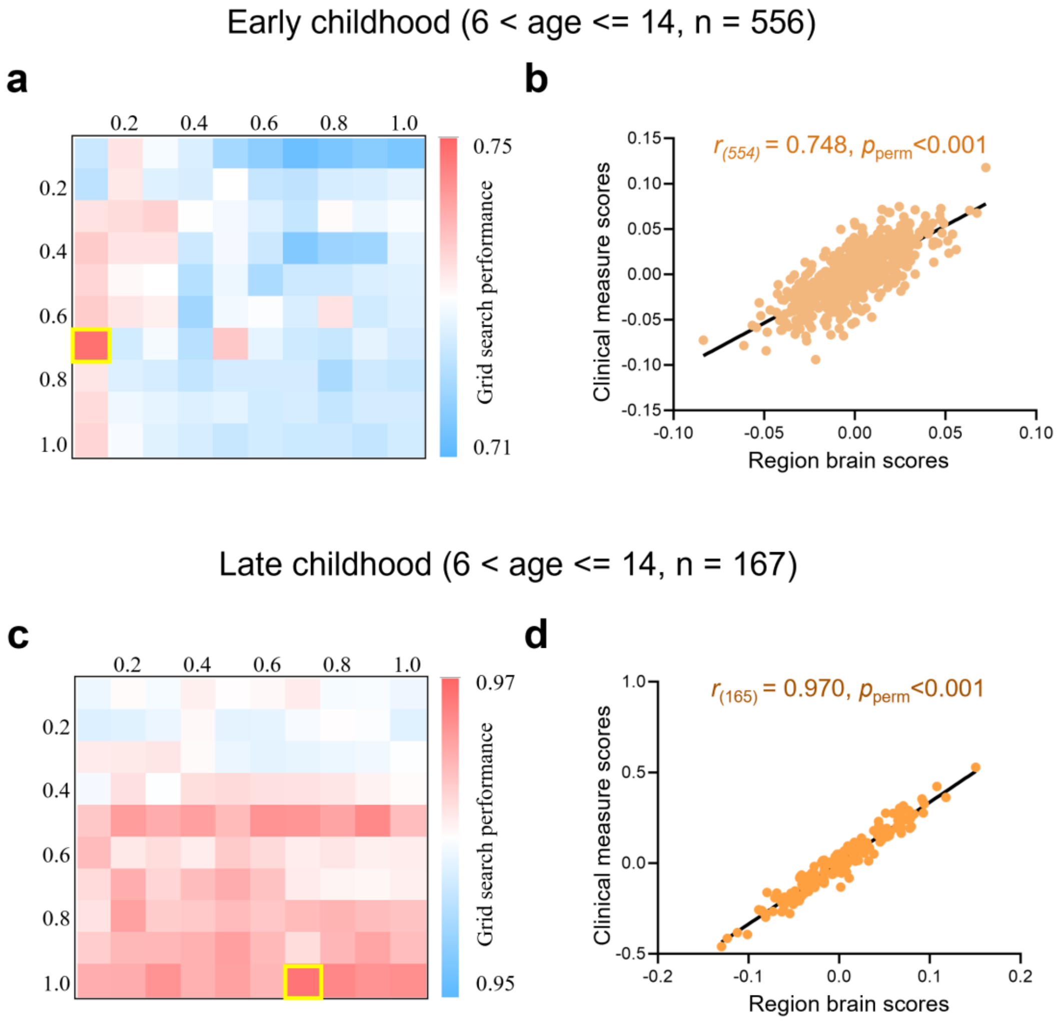


**Supplementary Fig. 3. Sparsity parameter search and identified first dimension of sparse canonical correlation analysis (sCCA). a,** Grid search for regularization parameters of sCCA in early childhood autism group, with a best regularization parameter of [0.7, 0.1]. **b,** Correlation between dynamic connectome lateralization strength (DCLS) and clinical measurement scores of the first linked dimensions in early childhood autism group. **c,** Grid search for regularization parameters of sCCA in late childhood autism group, with a best regularization parameter of [1.0, 0.7]. **d,** Correlation between DCLS and clinical measurement scores of the first linked dimensions on late childhood autism.

**Supplementary Table 1. Demographic characteristics and assessments information of SAED dataset.**

| Demographics | Early Childhood (0 < age ≤ 6 years) | | $p_{FDR-early}$ | Late Childhood (6 < age ≤ 14 years) | | $p_{FDR-late}$ |
|---|---|---|---|---|---|---|
| | Autism (n = 929) | TD (n = 147) | Autism vs. TD | Autism (n = 298) | TD (n = 179) | Autism vs. TD |
| Age, years, mean (SD) [range] | 3.67 (1.13) [1-6] | 3.94 (1.37) [7-14] | 0.008[a] | 7.93 (1.68) [7-14] | 8.72 (1.79) [7-14] | <0.001[a] |
| Sex ration, M/F | 768/161 | 68/79 | <0.001[b] | 245/53 | 70/109 | <0.001[b] |
| mean FD, mean (SD) [range] | 0.12 (0.07) [0.04-0.50] | 0.10 (0.05) [0.04-0.42] | 0.002[a] | 0.13 (0.08) [0.04-0.49] | 0.14 (0.08) [0.05-0.5] | 0.932[a] |
| ADOS-com, mean (SD) [range] | 5.74 (1.78) [0-10] | - | - | 5.40 (1.89) [1-10] | - | - |
| ADOS-soc, mean (SD) [range] | 10.01 (2.45) [1-14] | - | - | 9.52 (2.51) [1-14] | - | - |
| ADOS-rrb, mean (SD) [range] | 2.14 (1.59) [0-8] | - | - | 2.13 (1.60) [0-6] | - | - |
| ADOS-c+s, mean (SD) [range] | 15.79 (3.91) [1-24] | - | - | 14.94 (4.00) [6-24] | - | - |
| ADOS-total, mean (SD) [range] | 17.93 (4.90) [2-29] | - | - | 17.07 (4.97) [6-28] | - | - |
| SCQ-total, mean (SD) [range] | 17.16 (7.07) [0-32] | - | - | 14.90 (7.45) [0-32] | - | - |
| SEQ-hypoS, mean (SD) [range] | 5.33 (2.36) [0-12] | - | - | 4.81 (2.19) [0-10] | - | - |
| SEQ-hypoNS, mean (SD) [range] | 8.84 (4.96) [0-23] | - | - | 7.71 (5.54) [0-25] | - | - |
| SEQ-hypeS, mean (SD) [range] | 5.76 (3.20) [0-15] | - | - | 5.14 (3.38) [0-15] | - | - |
| SEQ-hypeNS, mean (SD) [range] | 6.65 (3.16) [0-16] | - | - | 6.57 (3.34) [0-16] | - | - |
| SEQ-total, mean (SD) [range] | 26.58 (10.73) [0-56] | - | - | 24.23 (10.88) [3-56] | - | - |
| SRS-awa, mean (SD) [range] | 11.24 (3.04) [3-22] | - | - | 11.17 (3.27) [4-22] | - | - |
| SRS-cog, mean (SD) [range] | 16.90 (4.72) [4-32] | - | - | 16.87 (4.63) [4-28] | - | - |
| SRS-com, mean (SD) [range] | 30.63 (9.59) [5-59] | - | - | 30.67 (9.18) [7-56] | - | - |
| SRS-mot, mean (SD) [range] | 13.79 (4.89) [1-28] | - | - | 14.01 (5.33) [4-26] | - | - |
| SRS-man, mean (SD) [range] | 12.70 (6.47) [0-34] | - | - | 13.19 (5.88) [0-28] | - | - |
| SRS-total, mean (SD) [range] | 85.25 (24.42) [25-168] | - | - | 85.92 (23.68) [26-153] | - | - |

SAED: Shanghai Autism Early Developmental Cohort; TD: typical development; SD: standard deviation; M: male; F: female; FD: framewise displacement; ADOS: Autism Diagnostic Observation Schedule; ADOS-com: ADOS communication; ADOS-soc: ADOS social interaction; ADOS-rrb: ADOS restricted and repetitive behaviors; ADOS-c+s: ADOS communication + social interaction; SCQ: Social Communication Questionnaire; SEQ: Sensory Experience Questionnaire; SEQ-hypoS: SEQ hypo-social; SEQ-hypoNS: SEQ hypo-nonsocial; SEQ-hypeS: SEQ hyper-social; SEQ-hypeNS: SEQ hyper-nonsocial; SRS: Social Responsiveness Scale; SRS-awa: SRS awareness; SRS-cog: SRS cognition; SRS-com: SRS communication; SRS-mot: SRS motivation; SRS-man: SRS mannerism;

[a] two-sample t test; [b] Chi-square test; all *p* values were corrected with FDR correction. 556 autistic individuals (59.8%) in early childhood group and 167 autistic individuals (56.0%) in late childhood group have complete assessments data.

**Supplementary Table 2. Demographic characteristics of validation dataset.**

| | **Autism (n = 56)** | **TD (n = 97)** | **Statistics** |
|---|---|---|---|
| **Site Name** | | | |
| ABIDE-I SDSU | 4 | 10 | |
| ABIDE-I YALE | 13 | 16 | |
| ABIDE-I LEUVEN_2 | 5 | 9 | |
| ABIDE-I KKI | 9 | 26 | |
| ABIDE-II KKI | 19 | 25 | |
| ABIDE-II UMIA | 7 | 11 | |
| **Demographics** | | | |
| Age, years, mean (SD) [range] | 10.77 (1.82) [7-14] | 10.75 (1.84) [7-14] | 0.944[a] |
| Sex ration, M/F | 43/13 | 70/27 | 0.002[b] |

TD: typical development; ABIDE: Autism Brain Imaging Data Exchange Cohort; SDSU: San Diego State University; YALE: Yale Child Study Center; LEUVEN: University of Leuven; KKI: Kennedy Krieger Institute; UMIA: University of Miami; SD: standard deviation; M: male, F: female.

[a], two-sample t test; [b], Chi-square test; all *p* values were corrected with FDR correction.

**Supplementary Table 3. Significant difference in dynamic connectome lateralization strength across the entire childhood period (0 < age ≤ 14 years) in discovery dataset.**

| **Gyrus** | **Region** | **Effect Size (Cohen's *d*)** | **t** | $p_{FDR}$ |
|---|---|---|---|---|
| Superior Frontal Gyrus | SFG_L_7_2 | 0.251 | 4.023 | 3.29E-04 |
| Superior Frontal Gyrus | SFG_R_7_2 | 0.279 | 4.470 | 6.45E-05 |
| Superior Frontal Gyrus | SFG_R_7_3 | 0.331 | 5.311 | 2.84E-06 |
| Superior Frontal Gyrus | SFG_L_7_4 | 0.319 | 5.120 | 5.65E-06 |
| Superior Frontal Gyrus | SFG_L_7_6 | 0.239 | 3.837 | 6.13E-04 |
| Superior Frontal Gyrus | SFG_L_7_7 | 0.288 | 4.616 | 3.36E-05 |
| Superior Frontal Gyrus | SFG_R_7_7 | 0.291 | 4.667 | 2.82E-05 |
| Middle Frontal Gyrus | MFG_L_7_1 | 0.356 | 5.707 | 4.22E-07 |
| Middle Frontal Gyrus | MFG_L_7_3 | 0.361 | 5.786 | 3.57E-07 |
| Middle Frontal Gyrus | MFG_L_7_4 | 0.321 | 5.150 | 5.16E-06 |
| Middle Frontal Gyrus | MFG_L_7_5 | 0.299 | 4.794 | 1.70E-05 |
| Middle Frontal Gyrus | MFG_L_7_6 | 0.309 | 4.953 | 1.05E-05 |
| Middle Frontal Gyrus | MFG_R_7_7 | 0.241 | 3.863 | 5.77E-04 |
| Inferior Frontal Gyrus | IFG_L_6_1 | 0.258 | 4.136 | 2.13E-04 |
| Inferior Frontal Gyrus | IFG_L_6_5 | 0.232 | 3.731 | 8.58E-04 |
| Inferior Frontal Gyrus | IFG_L_6_6 | 0.234 | 3.760 | 8.03E-04 |
| Orbital Gyrus | OrG_R_6_1 | 0.243 | 3.898 | 5.17E-04 |
| Orbital Gyrus | OrG_R_6_4 | 0.302 | 4.846 | 1.55E-05 |
| Orbital Gyrus | OrG_R_6_5 | 0.201 | 3.224 | 4.60E-03 |
| Orbital Gyrus | OrG_R_6_6 | 0.210 | 3.376 | 2.85E-03 |
| Precentral Gyrus | PrG_L_6_2 | 0.252 | 4.046 | 3.06E-04 |
| Precentral Gyrus | PrG_L_6_3 | 0.294 | 4.723 | 2.31E-05 |
| Precentral Gyrus | PrG_L_6_4 | 0.261 | 4.191 | 1.76E-04 |
| Precentral Gyrus | PrG_L_6_5 | 0.236 | 3.788 | 7.32E-04 |
| Precentral Gyrus | PrG_L_6_6 | 0.216 | 3.472 | 2.07E-03 |
| Paracentral Lobule | PCL_L_2_1 | 0.315 | 5.063 | 7.12E-06 |
| Superior Temporal Gyrus | STG_R_6_1 | 0.241 | 3.861 | 5.77E-04 |
| Superior Temporal Gyrus | STG_R_6_4 | 0.273 | 4.374 | 9.01E-05 |
| Superior Temporal Gyrus | STG_R_6_5 | 0.207 | 3.329 | 3.27E-03 |
| Superior Temporal Gyrus | STG_R_6_6 | 0.368 | 5.912 | 2.04E-07 |

| Gyrus | Region | Effect Size (Cohen's *d*) | t | $p_{FDR}$ |
|---|---|---|---|---|
| Middle Temporal Gyrus | MTG_R_4_1 | 0.306 | 4.915 | 1.21E-05 |
| Middle Temporal Gyrus | MTG_R_4_2 | 0.293 | 4.702 | 2.47E-05 |
| Middle Temporal Gyrus | MTG_L_4_3 | 0.305 | 4.899 | 1.25E-05 |
| Middle Temporal Gyrus | MTG_R_4_3 | 0.211 | 3.388 | 2.77E-03 |
| Middle Temporal Gyrus | MTG_R_4_4 | 0.387 | 6.216 | 8.04E-08 |
| Inferior Temporal Gyrus | ITG_L_7_2 | 0.248 | 3.979 | 3.87E-04 |
| Inferior Temporal Gyrus | ITG_L_7_5 | 0.331 | 5.308 | 2.84E-06 |
| Fusiform Gyrus | FuG_L_3_3 | 0.277 | 4.446 | 6.99E-05 |
| Parahippocampal Gyrus | PhG_L_6_6 | 0.204 | 3.272 | 3.95E-03 |
| posterior Superior Temporal Sulcus | pSTS_R_2_1 | 0.324 | 5.206 | 4.49E-06 |
| posterior Superior Temporal Sulcus | pSTS_L_2_2 | 0.220 | 3.533 | 1.74E-03 |
| posterior Superior Temporal Sulcus | pSTS_R_2_2 | 0.262 | 4.198 | 1.75E-04 |
| Superior Parietal Lobule | SPL_L_5_1 | 0.289 | 4.635 | 3.17E-05 |
| Superior Parietal Lobule | SPL_L_5_2 | 0.322 | 5.163 | 5.16E-06 |
| Superior Parietal Lobule | SPL_L_5_4 | 0.258 | 4.139 | 2.13E-04 |
| Inferior Parietal Lobule | IPL_L_6_1 | 0.354 | 5.676 | 4.49E-07 |
| Inferior Parietal Lobule | IPL_L_6_2 | 0.234 | 3.753 | 8.11E-04 |
| Inferior Parietal Lobule | IPL_R_6_2 | 0.239 | 3.837 | 6.13E-04 |
| Inferior Parietal Lobule | IPL_L_6_4 | 0.218 | 3.506 | 1.85E-03 |
| Inferior Parietal Lobule | IPL_R_6_5 | 0.373 | 5.993 | 1.73E-07 |
| Inferior Parietal Lobule | IPL_L_6_6 | 0.219 | 3.521 | 1.78E-03 |
| Precuneus | PCun_L_4_1 | 0.372 | 5.977 | 1.73E-07 |
| Precuneus | PCun_L_4_2 | 0.412 | 6.619 | 1.22E-08 |
| Precuneus | PCun_L_4_3 | 0.230 | 3.683 | 9.92E-04 |
| Precuneus | PCun_L_4_4 | 0.270 | 4.341 | 9.78E-05 |
| Precuneus | PCun_R_4_4 | 0.310 | 4.972 | 1.03E-05 |
| Postcentral Gyrus | PoG_R_4_1 | 0.209 | 3.346 | 3.13E-03 |
| Postcentral Gyrus | PoG_R_4_3 | 0.243 | 3.905 | 5.14E-04 |
| Postcentral Gyrus | PoG_L_4_4 | 0.301 | 4.837 | 1.55E-05 |

| **Gyrus** | **Region** | **Effect Size (Cohen's *d*)** | **t** | $p_{FDR}$ |
|---|---|---|---|---|
| Insular Gyrus | INS_L_6_3 | 0.270 | 4.331 | 9.94E-05 |
| Cingulate Gyrus | CG_R_7_1 | 0.356 | 5.719 | 4.22E-07 |
| Cingulate Gyrus | CG_L_7_3 | 0.299 | 4.797 | 1.70E-05 |
| Cingulate Gyrus | CG_R_7_4 | 0.230 | 3.692 | 9.77E-04 |
| Cingulate Gyrus | CG_L_7_5 | 0.274 | 4.400 | 8.36E-05 |
| Cingulate Gyrus | CG_L_7_6 | 0.299 | 4.803 | 1.70E-05 |
| lateral Occipital Cortex | LOcC_L_4_1 | 0.272 | 4.371 | 9.01E-05 |
| lateral Occipital Cortex | LOcC_L_4_2 | 0.232 | 3.729 | 8.58E-04 |
| lateral Occipital Cortex | LOcC_L_2_1 | 0.272 | 4.362 | 9.12E-05 |
| lateral Occipital Cortex | LOcC_L_2_2 | 0.309 | 4.967 | 1.03E-05 |

**Supplementary Table 4. Significant difference in dynamic connectome lateralization strength during the early childhood stage (0 < age ≤ 6 years) in discovery dataset.**

| Gyrus | Region | Effect Size (Cohen's *d*) | t | $p_{FDR}$ |
|---|---|---|---|---|
| Middle Frontal Gyrus | MFG_L_7_3 | 0.317 | 3.570 | 1.30E-02 |
| Middle Frontal Gyrus | MFG_L_7_4 | 0.348 | 3.922 | 1.15E-02 |
| Inferior Frontal Gyrus | IFG_L_6_1 | 0.276 | 3.108 | 2.50E-02 |
| Inferior Frontal Gyrus | IFG_L_6_2 | 0.305 | 3.434 | 1.52E-02 |
| Superior Temporal Gyrus | STG_R_6_4 | 0.361 | 4.062 | 1.15E-02 |
| Superior Temporal Gyrus | STG_R_6_6 | 0.323 | 3.643 | 1.16E-02 |
| Middle Temporal Gyrus | MTG_L_4_3 | 0.291 | 3.283 | 2.00E-02 |
| Middle Temporal Gyrus | MTG_R_4_4 | 0.282 | 3.181 | 2.18E-02 |
| Inferior Temporal Gyrus | ITG_L_7_5 | 0.276 | 3.111 | 2.50E-02 |
| Fusiform Gyrus | FuG_L_3_1 | 0.332 | 3.738 | 1.16E-02 |
| Fusiform Gyrus | FuG_L_3_2 | 0.286 | 3.222 | 2.07E-02 |
| Fusiform Gyrus | FuG_L_3_3 | 0.326 | 3.675 | 1.16E-02 |
| Inferior Parietal Lobule | IPL_L_6_1 | 0.325 | 3.661 | 1.16E-02 |
| Inferior Parietal Lobule | IPL_R_6_5 | 0.285 | 3.214 | 2.07E-02 |
| Precuneus | PCun_R_4_4 | 0.289 | 3.251 | 2.07E-02 |
| Insular Gyrus | INS_L_6_3 | 0.305 | 3.440 | 1.52E-02 |
| Cingulate Gyrus | CG_R_7_1 | 0.293 | 3.303 | 2.00E-02 |
| lateral Occipital Cortex | LOcC_L_4_4 | 0.293 | 3.299 | 2.00E-02 |
| Amygdala | Amyg_L_2_1 | 0.273 | 3.081 | 2.61E-02 |
| Amygdala | Amyg_L_2_2 | 0.314 | 3.537 | 1.30E-02 |

**Supplementary Table 5. Significant difference in dynamic connectome lateralization strength during late childhood stage (6 < age ≤ 14 years) in discovery dataset.**

| Gyrus | Region | Effect Size (Cohen's *d*) | t | $p_{FDR}$ |
|---|---|---|---|---|
| Superior Frontal Gyrus | SFG_R_7_2 | 0.269 | 2.842 | 2.95E-02 |
| Superior Frontal Gyrus | SFG_R_7_3 | 0.357 | 3.777 | 3.67E-03 |
| Superior Frontal Gyrus | SFG_L_7_4 | 0.374 | 3.950 | 2.58E-03 |
| Superior Frontal Gyrus | SFG_L_7_7 | 0.271 | 2.864 | 2.90E-02 |
| Middle Frontal Gyrus | MFG_L_7_1 | 0.325 | 3.440 | 7.08E-03 |
| Middle Frontal Gyrus | MFG_L_7_3 | 0.336 | 3.558 | 5.32E-03 |
| Middle Frontal Gyrus | MFG_L_7_4 | 0.273 | 2.892 | 2.82E-02 |
| Middle Frontal Gyrus | MFG_L_7_5 | 0.261 | 2.765 | 3.38E-02 |
| Middle Frontal Gyrus | MFG_L_7_6 | 0.386 | 4.084 | 2.13E-03 |
| Inferior Frontal Gyrus | IFG_L_6_1 | 0.279 | 2.946 | 2.51E-02 |
| Inferior Frontal Gyrus | IFG_L_6_5 | 0.353 | 3.732 | 3.87E-03 |
| Inferior Frontal Gyrus | IFG_L_6_6 | 0.341 | 3.606 | 4.95E-03 |
| Precentral Gyrus | PrG_L_6_2 | 0.265 | 2.801 | 3.11E-02 |
| Precentral Gyrus | PrG_L_6_3 | 0.342 | 3.620 | 4.95E-03 |
| Precentral Gyrus | PrG_L_6_4 | 0.352 | 3.723 | 3.87E-03 |
| Precentral Gyrus | PrG_L_6_5 | 0.359 | 3.793 | 3.67E-03 |
| Paracentral Lobule | PCL_L_2_1 | 0.455 | 4.809 | 2.51E-04 |
| Superior Temporal Gyrus | STG_R_6_1 | 0.373 | 3.947 | 2.58E-03 |
| Superior Temporal Gyrus | STG_R_6_6 | 0.340 | 3.592 | 4.95E-03 |
| Middle Temporal Gyrus | MTG_R_4_1 | 0.287 | 3.036 | 2.04E-02 |
| Middle Temporal Gyrus | MTG_R_4_2 | 0.319 | 3.378 | 7.95E-03 |
| Middle Temporal Gyrus | MTG_L_4_3 | 0.326 | 3.444 | 7.08E-03 |
| Middle Temporal Gyrus | MTG_R_4_4 | 0.409 | 4.329 | 8.98E-04 |
| Inferior Temporal Gyrus | ITG_L_7_2 | 0.286 | 3.021 | 2.04E-02 |
| Inferior Temporal Gyrus | ITG_L_7_5 | 0.333 | 3.524 | 5.73E-03 |
| Fusiform Gyrus | FuG_L_3_3 | 0.259 | 2.738 | 3.59E-02 |
| posterior Superior Temporal Sulcus | pSTS_R_2_1 | 0.363 | 3.833 | 3.53E-03 |
| posterior Superior Temporal Sulcus | pSTS_L_2_2 | 0.275 | 2.905 | 2.78E-02 |
| posterior Superior Temporal Sulcus | pSTS_R_2_2 | 0.286 | 3.027 | 2.04E-02 |
| Superior Parietal Lobule | SPL_L_5_1 | 0.322 | 3.403 | 7.73E-03 |

| Gyrus | Region | Effect Size (Cohen's $d$) | t | $p_{FDR}$ |
|---|---|---|---|---|
| Superior Parietal Lobule | SPL_L_5_2 | 0.272 | 2.878 | 2.86E-02 |
| Superior Parietal Lobule | SPL_L_5_4 | 0.350 | 3.697 | 3.99E-03 |
| Inferior Parietal Lobule | IPL_L_6_1 | 0.372 | 3.938 | 2.58E-03 |
| Inferior Parietal Lobule | IPL_R_6_5 | 0.409 | 4.330 | 8.98E-04 |
| Inferior Parietal Lobule | IPL_L_6_6 | 0.258 | 2.723 | 3.67E-02 |
| Precuneus | PCun_L_4_1 | 0.311 | 3.285 | 1.04E-02 |
| Precuneus | PCun_L_4_2 | 0.476 | 5.028 | 1.73E-04 |
| Postcentral Gyrus | PoG_R_4_3 | 0.266 | 2.818 | 3.10E-02 |
| Postcentral Gyrus | PoG_L_4_4 | 0.430 | 4.548 | 5.66E-04 |
| Cingulate Gyrus | CG_R_7_1 | 0.319 | 3.372 | 7.95E-03 |
| Cingulate Gyrus | CG_L_7_3 | 0.304 | 3.218 | 1.25E-02 |
| Cingulate Gyrus | CG_L_7_5 | 0.248 | 2.618 | 4.88E-02 |
| MedioVentral Occipital Cortex | MVOcC_L_5_5 | 0.265 | 2.802 | 3.11E-02 |
| lateral Occipital Cortex | LOcC_L_4_1 | 0.270 | 2.856 | 2.90E-02 |
| lateral Occipital Cortex | LOcC_L_2_1 | 0.300 | 3.173 | 1.41E-02 |
| lateral Occipital Cortex | LOcC_L_2_2 | 0.292 | 3.092 | 1.78E-02 |

**Supplementary Table 6. Significant log variability ratio of dynamic connectome lateralization strength between autism and TD individuals during the late childhood stage (6 < age ≤ 14 years) in discovery dataset.**

| Gyrus | Region | lnVR | 95%CI | *p* |
|---|---|---|---|---|
| Inferior Frontal Gyrus | IFG_L_6_5 | 0.331 | [0.068, 0.595] | 0.014 |
| Orbital Gyrus | OrG_R_6_1 | -0.337 | [-0.600, -0.073] | 0.012 |
| Superior Temporal Gyrus | STG_L_6_2 | 0.299 | [0.036, 0.562] | 0.026 |
| Superior Temporal Gyrus | STG_L_6_6 | 0.300 | [0.038, 0.565] | 0.025 |
| Middle Temporal Gyrus | MTG_L_4_1 | 0.267 | [0.004, 0.531] | 0.047 |
| Middle Temporal Gyrus | MTG_L_4_4 | 0.264 | [0.001, 0.528] | 0.049 |
| Inferior Temporal Gyrus | ITG_L_7_4 | 0.367 | [0.103, 0.630] | 0.006 |
| Inferior Temporal Gyrus | ITG_L_7_7 | 0.277 | [0.013, 0.540] | 0.040 |
| Parahippocampal Gyrus | PhG_L_6_1 | 0.292 | [0.028, 0.555] | 0.030 |
| Parahippocampal Gyrus | PhG_L_6_3 | 0.279 | [0.015, 0.542] | 0.038 |
| Parahippocampal Gyrus | PhG_L_6_5 | 0.414 | [0.153, 0.680] | 0.002 |
| posterior Superior Temporal Sulcus | pSTS_L_2_1 | 0.287 | [0.024, 0.551] | 0.033 |
| Insular Gyrus | INS_R_6_2 | -0.268 | [-0.531, -0.004] | 0.047 |
| Insular Gyrus | INS_L_6_3 | 0.319 | [0.056, 0.583] | 0.017 |
| Insular Gyrus | INS_R_6_4 | -0.279 | [-0.542, -0.015] | 0.038 |
| Insular Gyrus | INS_L_6_6 | 0.303 | [0.040, 0.567] | 0.024 |
| MedioVentral Occipital Cortex | MVOcC_R_5_1 | -0.301 | [-0.565, -0.038] | 0.025 |
| MedioVentral Occipital Cortex | MVOcC_L_5_3 | -0.273 | [-0.536, -0.009] | 0.043 |
| lateral Occipital Cortex | LOcC_L_4_3 | -0.271 | [-0.533, -0.007] | 0.044 |
| Amygdala | Amyg_L_2_2 | 0.350 | [0.086, 0.613] | 0.009 |

**Supplementary Table 7. Significant regional brain loadings of the first canonical variate during the early childhood stage (0 < age ≤ 6 years) in discovery dataset.**

| Gyrus | Region | Brain loading | $p_{perm}$_FDR |
|---|---|---|---|
| Superior Frontal Gyrus | SFG_L_7_3 | 0.181 | <1.0E-04 |
| Superior Frontal Gyrus | SFG_R_7_4 | 0.133 | 2.13E-02 |
| Inferior Frontal Gyrus | IFG_L_6_3 | 0.133 | 2.13E-02 |
| Orbital Gyrus | OrG_R_6_6 | 0.151 | 6.15E-03 |
| Superior Temporal Gyrus | STG_L_6_1 | 0.120 | 2.13E-02 |
| Superior Temporal Gyrus | STG_R_6_5 | 0.148 | 9.84E-03 |
| Superior Temporal Gyrus | MTG_R_4_3 | 0.136 | 1.76E-02 |
| Inferior Temporal Gyrus | ITG_L_7_1 | 0.120 | 3.33E-02 |
| Inferior Temporal Gyrus | ITG_L_7_6 | 0.137 | 1.76E-02 |
| Inferior Temporal Gyrus | ITG_L_7_7 | 0.113 | 3.94E-02 |
| Parahippocampal Gyrus | PhG_L_6_5 | 0.116 | 3.63E-02 |
| Precuneus | PCun_R_4_3 | 0.138 | 1.91E-02 |
| Postcentral Gyrus | PoG_R_4_1 | 0.124 | 2.15E-02 |
| Postcentral Gyrus | PoG_L_4_3 | 0.128 | 2.13E-02 |
| Insular Gyrus | INS_R_6_2 | 0.138 | 6.15E-03 |
| MedioVentral Occipital Cortex | MVOcC_R_5_5 | 0.128 | 2.13E-02 |
| lateral Occipital Cortex | LOcC_R_2_1 | 0.143 | 1.91E-02 |
| Amygdala | Amyg_R_2_1 | 0.129 | 2.13E-02 |
| Amygdala | Amyg_R_2_2 | 0.186 | <1.0E-04 |
| Basal Ganglia | BG_L_6_1 | 0.115 | 3.63E-02 |

**Supplementary Table 8. Significant regional brain loadings of the first canonical variate during the late childhood stage (6 < age ≤ 14 years) in discovery dataset.**

| Gyrus | Region | Brain loading | $p_{perm\_FDR}$ |
|---|---|---|---|
| Superior Frontal Gyrus | SFG_L_7_1 | 0.243 | 2.46E-02 |
| Superior Frontal Gyrus | SFG_L_7_2 | 0.229 | 2.68E-02 |
| Superior Frontal Gyrus | SFG_L_7_3 | 0.225 | 3.11E-02 |
| Superior Frontal Gyrus | SFG_L_7_6 | 0.215 | 3.11E-02 |
| Superior Frontal Gyrus | SFG_L_7_7 | 0.189 | 4.84E-02 |
| Middle Frontal Gyrus | MFG_L_7_1 | 0.296 | <1.0E-04 |
| Middle Frontal Gyrus | MFG_L_7_3 | 0.200 | 4.13E-02 |
| Middle Frontal Gyrus | MFG_L_7_5 | 0.316 | <1.0E-04 |
| Middle Frontal Gyrus | MFG_L_7_6 | 0.214 | 3.11E-02 |
| Inferior Frontal Gyrus | IFG_L_6_2 | 0.191 | 4.67E-02 |
| Inferior Frontal Gyrus | IFG_R_6_2 | 0.207 | 4.13E-02 |
| Inferior Frontal Gyrus | IFG_R_6_3 | 0.191 | 4.84E-02 |
| Inferior Frontal Gyrus | IFG_L_6_4 | 0.253 | 2.46E-02 |
| Inferior Frontal Gyrus | IFG_R_6_4 | 0.196 | 4.48E-02 |
| Inferior Frontal Gyrus | IFG_L_6_6 | 0.206 | 4.13E-02 |
| Inferior Frontal Gyrus | IFG_R_6_6 | 0.213 | 3.11E-02 |
| Orbital Gyrus | OrG_L_6_2 | 0.204 | 4.13E-02 |
| Orbital Gyrus | OrG_L_6_6 | 0.244 | 2.68E-02 |
| Precentral Gyrus | PrG_R_6_6 | 0.287 | 4.92E-03 |
| Superior Temporal Gyrus | STG_R_6_3 | 0.194 | 4.67E-02 |
| Superior Temporal Gyrus | STG_R_6_4 | 0.197 | 4.48E-02 |
| Middle Temporal Gyrus | MTG_L_4_1 | 0.221 | 3.11E-02 |
| Inferior Temporal Gyrus | ITG_R_7_2 | 0.220 | 3.11E-02 |
| posterior Superior Temporal Sulcus | pSTS_L_2_1 | 0.232 | 2.87E-02 |
| posterior Superior Temporal Sulcus | pSTS_L_2_2 | 0.223 | 3.11E-02 |
| Inferior Parietal Lobule | IPL_R_6_3 | 0.196 | 4.13E-02 |
| Inferior Parietal Lobule | IPL_L_6_5 | 0.294 | <1.0E-04 |
| Postcentral Gyrus | PoG_R_4_2 | 0.231 | 2.11E-02 |
| Insular Gyrus | INS_L_6_1 | 0.200 | 4.48E-02 |
| Amygdala | Amyg_R_2_1 | 0.340 | <1.0E-04 |

| Gyrus | Region | Brain loading | $p_{perm}$_FDR |
|---|---|---|---|
| Amygdala | Amyg_R_2_2 | 0.209 | 4.13E-02 |
| Hippocampus | Hipp_R_2_1 | 0.268 | 8.20E-03 |

**Supplementary Table 9. Person's correlation between each neurotransmitter receptor and DCLS abnormalities in early childhood autistic individuals**

| Receptor/ transporter | left-hemispheric | | right-hemispheric | |
|---|---|---|---|---|
| | ***r*** **value** | $p_{spin\text{-}FDR}$ | ***r*** **value** | $p_{spin\text{-}FDR}$ |
| 5-$HT_{1A}$ | 0.296 | **1.90E-02** | 0.148 | 1.91E-01 |
| 5-$HT_{1B}$ | 0.093 | 4.08E-01 | 0.199 | 9.07E-02 |
| 5-$HT_2$ | 0.145 | 2.03E-01 | 0.332 | **3.80E-03** |
| 5-$HT_4$ | -0.047 | 6.33E-01 | 0.195 | 9.07E-02 |
| 5-$HT_6$ | 0.063 | 6.15E-01 | 0.081 | 4.48E-01 |
| 5-HTT | -0.219 | **4.89E-02** | -0.227 | 7.73E-02 |
| $\alpha_4\beta_2$ | -0.272 | **2.47E-02** | -0.194 | 9.07E-02 |
| $CB_1$ | 0.218 | **4.89E-02** | 0.167 | 1.39E-01 |
| $D_1$ | -0.057 | 6.15E-01 | 0.045 | 7.00E-01 |
| $D_2$ | -0.130 | 2.45E-01 | -0.109 | 3.14E-01 |
| DAT | -0.052 | 6.33E-01 | -0.128 | 2.81E-01 |
| $GABA_A$ | 0.158 | 1.72E-01 | 0.202 | 9.07E-02 |
| $H_3$ | -0.017 | 8.59E-01 | 0.001 | 9.97E-01 |
| $M_1$ | 0.231 | **4.89E-02** | 0.167 | 1.39E-01 |
| MOR | -0.105 | 3.64E-01 | 0.000 | 9.97E-01 |
| NET | -0.214 | **4.89E-02** | -0.118 | 2.86E-01 |
| NMDA | -0.179 | 1.17E-01 | -0.119 | 2.86E-01 |
| VAChT | -0.140 | 2.20E-01 | -0.089 | 4.13E-01 |
| $mGluR_5$ | 0.235 | **4.81E-02** | 0.326 | **3.80E-03** |

**Supplementary Table 10. Person's correlation between each neurotransmitter receptor and DCLS abnormalities in late childhood autistic individuals**

| Receptor/ transporter | left-hemispheric | | right-hemispheric | |
|---|---|---|---|---|
| | ***r*** **value** | $p_{spin\text{-}FDR}$ | ***r*** **value** | $p_{spin\text{-}FDR}$ |
| 5-$HT_{1A}$ | -0.036 | 7.27E-01 | 0.473 | **<1.0E-04** |
| 5-$HT_{1B}$ | 0.415 | **<1.0E-04** | 0.063 | 4.90E-01 |
| 5-$HT_{2}$ | 0.290 | **1.90E-03** | 0.497 | **<1.0E-04** |
| 5-$HT_{4}$ | -0.053 | 6.19E-01 | -0.070 | 4.57E-01 |
| 5-$HT_{6}$ | 0.146 | 1.41E-01 | -0.174 | 6.49E-02 |
| 5-HTT | -0.325 | **<1.0E-04** | -0.452 | **<1.0E-04** |
| $\alpha_4\beta_2$ | -0.110 | 2.83E-01 | -0.339 | **4.75E-04** |
| $CB_1$ | 0.074 | 4.95E-01 | 0.198 | **3.09E-02** |
| $D_1$ | -0.178 | 7.74E-02 | -0.238 | **1.14E-02** |
| $D_2$ | -0.267 | **9.50E-03** | -0.365 | **<1.0E-04** |
| DAT | -0.278 | **5.97E-03** | -0.361 | **<1.0E-04** |
| $GABA_A$ | 0.321 | **1.90E-03** | 0.226 | **1.67E-02** |
| $H_3$ | 0.026 | 7.86E-01 | -0.347 | **<1.0E-04** |
| $M_1$ | 0.188 | 6.46E-02 | 0.099 | 3.14E-01 |
| MOR | -0.159 | 1.20E-01 | -0.118 | 2.31E-01 |
| NET | 0.296 | **1.90E-03** | -0.279 | **3.80E-03** |
| NMDA | -0.241 | **1.56E-02** | -0.306 | **1.27E-03** |
| VAChT | -0.158 | 1.16E-01 | -0.442 | **<1.0E-04** |
| $mGluR_5$ | 0.341 | **1.27E-03** | 0.233 | **1.14E-02** |

**Supplementary Table 11. Scan parameters of the sMRI data from each site of SAED and ABIDE cohort.**

| Site | Scanner | TR (ms) | TE (ms) | FA (°) | Thickness (mm) |
|---|---|---|---|---|---|
| SAED | Siemens Verio 3.0T | 2300 | 2.28 | 8 | 1 |
| SAED | Philips Achieva 3T | 6.12 | 2.81 | 8 | 1 |
| ABIDE-I SDSU | GE 3T MR750 | - | Min Full | 8 | 1 |
| ABIDE-I YALE | Siemens Trio 3T | 1230 | 1.73 | 9 | 1 |
| ABIDE-I LEUVEN_2 | Philips 3T | shortest | 4.6 | 8 | 1.2 |
| ABIDE-I KKI | Philips Achieva 3T | 8.0 | 3.7 | 8 | 1 |
| ABIDE-II KKI | Philips 3T | 8.0 | 3.7 | 8 | 1 |
| ABIDE-II UMIA | GE Healthcare 3T | - | - | 12 | 1 |

SAED: Shanghai Autism Early Developmental Cohort; ABIDE: Autism Brain Imaging Data Exchange Cohort; TR: repetition time; TE: echo time; FA: flip angle; TD: typical development; ABIDE: Autism Brain Imaging Data Exchange; SDSU: San Diego State University; YALE: Yale Child Study Center; LEUVEN: University of Leuven; KKI: Kennedy Krieger Institute; UMIA: University of Miami.

**Supplementary Table 12. Scan parameters of the rsfMRI data from each site of SAED and ABIDE cohort.**

| Site | Scanner | TR (ms) | TE (ms) | FA (°) | Time Points | Thickness (mm) |
|---|---|---|---|---|---|---|
| SAED | Siemens Verio 3.0T | 2000 | 30 | 90 | 200 | 4 |
| SAED | Philips Achieva 3T | 1400 | 25 | 65 | 260 | 3 |
| ABIDE-I SDSU | GE 3T MR750 | 2000 | 30 | 45 | 180 | 3.4 |
| ABIDE-I YALE | Siemens Trio 3T | 2000 | 25 | 60 | 200 | 4 |
| ABIDE-I LEUVEN_2 | Philips 3T | 1667 | 33 | 90 | 250 | 4 |
| ABIDE-I KKI | Philips Achieva 3T | 2500 | 30 | 75 | 156 | 3 |
| ABIDE-II KKI | Philips 3T | 2500 | 30 | 90 | 156 | 2.7 |
| ABIDE-II UMIA | GE Healthcare 3T | 2000 | 30 | 75 | 295 | 3.4 |

SAED: Shanghai Autism Early Developmental Cohort; ABIDE: Autism Brain Imaging Data Exchange Cohort; TR: repetition time; TE: echo time; FA: flip angle; TD: typical development; ABIDE: Autism Brain Imaging Data Exchange; SDSU: San Diego State University; YALE: Yale Child Study Center; LEUVEN: University of Leuven; KKI: Kennedy Krieger Institute; UMIA: University of Miami.

**Supplementary Table 13. Information of PET images for each neurotransmitter receptors and transporters.**

| Receptor/ transporter | Neurotransmitter | Tracer | Measure | n | Age | Reference |
|---|---|---|---|---|---|---|
| 5-HT$_{1A}$ | Serotonin | [$^{11}$C]WAY-100635 | BP$_{ND}$ | 35 | 26.3 ± 5.2 | Savli et al.[5] |
| 5-HT$_{1B}$ | Serotonin | [$^{11}$C]P943 | BP$_{ND}$ | 23 | 28.7 ± 7.0 | Savli et al.[5] |
| 5-HT$_{2}$ | Serotonin | [$^{11}$C]Cimbi-36 | B$_{max}$ | 29 | 22.6 ± 2.7 | Beliveau et al.[6] |
| 5-HT$_{4}$ | Serotonin | [$^{11}$C]SB207145 | B$_{max}$ | 59 | 25.9 ± 5.3 | Beliveau et al.[6] |
| 5-HT$_{6}$ | Serotonin | [$^{11}$C]GSK215083 | BP$_{ND}$ | 30 | 36.6 ± 9.0 | Radhakrishnan et al.[7,8] |
| 5-HTT | Serotonin | [$^{11}$C]DASB | B$_{max}$ | 100 | 25.1 ± 5.8 | Beliveau et al.[6] |
| $\alpha_4\beta_2$ | Acetylcholine | [$^{18}$F]Flubatine | V$_{T}$ | 30 | 33.5 ± 10.7 | Hillmer et al.[9,10] |
| CB$_{1}$ | Cannabinoid | [$^{11}$C]OMAR | V$_{T}$ | 77 | 30.0 ± 8.9 | Normandin et al.[11–14] |
| D$_{1}$ | Dopamine | [$^{11}$C]SCH23390 | BP$_{ND}$ | 13 | 33.0 ± 13.0 | Kaller et al.[15] |
| D$_{2}$ | Dopamine | [$^{11}$C]FLB-457 | BP$_{ND}$ | 55 | 32.5 ± 9.7 | Sandiego et al.[16–20] |
| DAT | Dopamine | [$^{123}$I]-FP-CIT | SUVR | 65 | 61.0 ± 11.0 | Dukart et al.[21] |
| GABA$_{A}$ | γ-aminobutyric | [$^{11}$C]Flumazenil | Bmax | 16 | 26.6 ± 8.0 | Nørgaard et al.[22] |
| H$_{3}$ | Histamine | [$^{11}$C]GSK189254 | V$_{T}$ | 8 | 31.7 ± 9.0 | Gallezot et al.[23] |
| M$_{1}$ | Acetylcholine | [$^{11}$C]LSN3172176 | BP$_{ND}$ | 24 | 40.5 ± 11.7 | Naganawa et al.[24] |
| MOR | Opioid | [$^{11}$C]Carfentanil | BP$_{ND}$ | 204 | 32.3 ± 10.8 | Kantonen et al.[25] |
| NET | Norepinephrine | [$^{11}$C]MRB | BP$_{ND}$ | 77 | 33.4 ± 9.2 | Ding et al.[26–29] |
| NMDA | Glutamate | [$^{18}$F]GE-179 | V$_{T}$ | 29 | 40.9 ± 12.7 | Galovic et al.[30–32] |
| VAChT | Acetylcholine | [$^{18}$F]FEOBV | SUVR | 18 | 66.8 ± 6.8 | Aghourian et al.[33] |
| mGluR$_{5}$ | Glutamate | [$^{11}$C]ABP688 | BP$_{ND}$ | 73 | 19.9 ± 3.04 | Smart et al.[34] |

BP$_{ND}$, non-displaceable binding potential; V$_{T}$, tracer distribution volume; B$_{max}$, density (pmol ml$^{-1}$) converted from binding potential (5-HT) or distributional volume (GABA) using autoradiography-derived densities; SUVR, standard uptake value ratio.
